# A Useful Guide to Lectin Binding: Machine-Learning Directed Annotation of 57 Unique Lectin Specificities

**DOI:** 10.1101/2021.08.31.458439

**Authors:** Daniel Bojar, Lawrence Meche, Guanmin Meng, William Eng, David F. Smith, Richard D. Cummings, Lara K. Mahal

## Abstract

Glycans are critical to every facet of biology and medicine, from viral infections to embryogenesis. Tools to study glycans are rapidly evolving, however the majority of our knowledge is deeply dependent on binding by glycan binding proteins (e.g., lectins). The specificities of lectins, which are often naturally isolated proteins, have not been well-defined, making it difficult to leverage their full potential for glycan analysis. Herein, we use glycan microarray analysis of 116 commercially available lectins, including different preparations of the same lectin, to extract the specific glycan features required for lectin binding. Data was obtained using the Consortium for Functional Glycomics microarray (CFG v5.0) containing 611 glycans. We use a combination of machine learning algorithms to define lectin specificity, mapping inputs (glycan sequences) to outputs (lectin-glycan binding) for a large-scale evaluation of lectin-glycan binding behaviours. Our motif analysis was performed by integrating 68 manually defined glycan features with systematic probing of computational rules for significant binding motifs using mono- and disaccharides- and linkages. Using a combination of machine learning and manual annotation of the data, we created a detailed interpretation of glycan-binding specificity for 57 unique lectins, categorized by their major binding motifs: mannose, complex-type *N*-glycan, *O*-glycan, fucose, sialic acid and sulfate, GlcNAc and chitin, Gal and LacNAc, and GalNAc. Our work provides fresh insights into the complex binding features of commercially available lectins in current use, providing a critical guide to these important reagents.

## INTRODUCTION

Carbohydrates (i.e., glycans) are involved in every facet of life from the cell walls of bacteria to the signals that start inflammatory cascades in humans [2]. While our understanding of other biomolecules, such as DNA and RNA, has expanded exponentially due to the advent of new analytical technologies, glycans have remained understudied due to the lack of convenient analytical tools. Nature solved the problem of glycan identification by designing non-catalytic glycan binding proteins called lectins, which recognize well-defined epitopes within a glycan. Lectins, which are often isolated from plants, have long been used as tools for glycan analysis [3]. Indeed our earliest understanding of blood group antigens comes from agglutination studies using lectins to determine blood type [4]. More modern methods that leverage lectins as analytical tools include lectin histology and enzyme-linked lectin assays (ELLA) [4]. Lectin microarrays, in which a panel of lectins (10s to >100) and other carbohydrate-binding probes (e.g., antibodies) are printed on a solid support, are now routinely used in glycomics [5–7]. This high-throughput method has identified glycans involved in a variety of systems from melanoma metastasis to viral host-response [8–10]. Although lectins are useful tools, they have suffered from a lack of detailed definition of their binding requirements, which hampers their use analytically. Traditional methods of defining lectin specificity have involved inhibition assays with monosaccharides, binding assays with a limited set of potential ligands, or in a few cases, crystal structures with disaccharide or monosaccharide binder. Few analyses have examined the same lectin from different sources to identify the core binding motifs. To leverage the full potential of lectins for glycan analysis, a more rigorous characterization of lectin specificity must be obtained.

The best approach to define the specificity of a lectin would be to test it against an extensive library of glycan epitopes. In this study, we characterize the glycan binding specificities of 116 commercially-available lectins using version 5.0 of the Consortium for Functional Glycomics (CFG) glycan microarray and a combination of machine learning and manual annotation [11].

Connecting binding events with glycan substructures or motifs has attracted considerable interest for the purpose of identifying glycan receptors and has led to the development of multiple algorithms including frequent subtree mining [12, 13] and motif based approaches[14, 15]. Yet the high diversity and non-linearity of glycans has stymied the large-scale evaluation of subtle, interpretable binding motifs in glycan array data to date. To overcome this, we have leveraged the recent introduction of machine learning into glycobiology [16]. By mapping inputs (glycan sequences) to outputs (lectin-glycan binding), machine learning algorithms can ascertain the most important features (i.e., sequence motifs) that predict lectin-glycan binding. Importantly, this is performed on a scale that is vastly larger than manual annotation and also enables the analysis of highly complex feature combinations to obtain insights into more subtle influences of the co-occurrence of glycan motifs.

We further engaged in feature engineering by combining hand-crafted features that are domain-relevant (e.g., the presence of Lewis A or terminal sialosides) with systematic probing for all observed sequence motifs of various lengths. This procedure improves the interpretability of the resulting machine learning models, which have traditionally been labelled as uninterpretable “black boxes”. We then used these features, our machine learning models, and iterative manual annotation to establish logical, interpretable rules for each lectin that best explained lectin-glycan binding behavior to further facilitate “white box” machine learning and extract lectin binding specificities. Our analysis, which includes preparations of the same lectin from several sources, provides detailed annotation of 57 unique commercially available lectins. The results of our work give a clear view into the binding profiles of these carbohydrate-binding reagents and demonstrates that they are selective in their epitope binding. Overall, our work provides a useful guide to anyone analyzing glycans using this reagent class and sets the stage for more advanced interpretation of studies using lectins.

## RESULTS AND DISCUSSION

### Study Design

The glycan binding specificities of commercially available lectins from various sources (116 total) were analyzed on the CFG glycan microarray version 5.0. (CFGv5). This array contains 611 glycan structures, including those representing both *N-* and *O-* glycans (**Figure 1).** *N-*glycans are the most heavily studied glycan cohort. They are attached to asparagine and follow a well-defined biosynthetic pathway. Of the epitopes on CFGv5, 22.3% represent *N-*linked glycans, with the exception of hybrid *N*-linked sugars, which are not on this array. Another 18.5% represent *O*-linked glycans, which do not have a conserved core and are attached to either serines or threonines. The *O-*glycan subset focuses on short *O-*glycans of various core structures, with little representation of more elaborate structures. The remaining array epitopes are either terminal or fragmented glycans, which can appear on either *N-* or *O-*linked carbohydrates, or select glycolipid structures (59%). All glycans are attached to the slides via NHS-coupling to either aliphatic amine linkers of varying lengths or the amino termini of amino acids (serine, threonine, asparagine, glycine) [4][17]. In this study, 103 of the 611 glycans on the array did not bind to any of the commercial lectins tested, indicating that either there were no binders for the epitope or, in some instances, there may be a misprint on the array (for an annotated list of glycans see **Table S1**).

**FIGURE 1.**
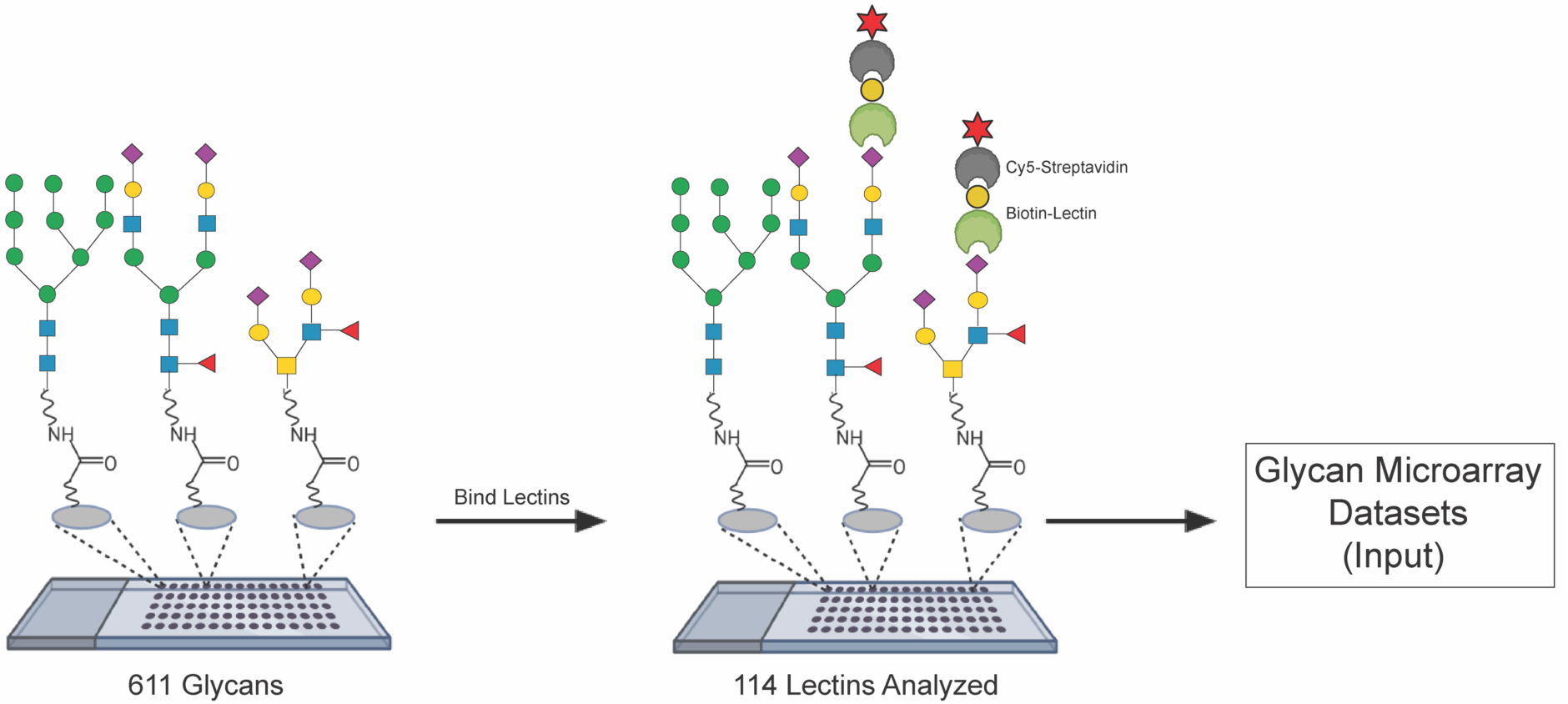
Glycan microarray datasets were generated with the Consortium for Functional Glycomics glycan microarray version 5 (CFGv5) containing 611 unique glycans. Biotinylated lectins (116 total) were incubated with the arrays at varying concentrations, followed by incubation with Cy5- Streptavidin. Slides were processed, scanned and data was extracted. For each array, the average fluorescence for each glycan (n= 4 spots) was obtained. Array datasets were then used as input for machine learning and Z-score analysis.

The 116 commercially available lectins analyzed included multiple lots of the same lectin from a variety of sources (**Table S2)**. Where available, the biotinylated versions of lectins were purchased. Lectins that were available only in an unmodified form were biotinylated for analysis using standard protocols (AOL, CF, PA-IL, PAA, SVAM, TJA-I, TJA-II). In general, lectins were incubated with the array at 3-6 concentrations ranging from 0.1-100 μg/mL and binding was detected via Cy5-labeled streptavidin (**Figure 1**) [11]. Analyzing several concentrations of a lectin allows for better separation between the strong and weak binders through sampling of a greater cross-section of the binding interactions. For each array, fluorescence data was extracted for the glycans, which were represented by 6 spots per glycan. The average fluorescence of four spots, excluding the high and low values, was obtained for each glycan and used in our data analysis. Lectins that showed no binding ≥1000 RFUs at the concentrations tested or that had data for only a single concentration of that lectin with signal ≥1000 RFU were excluded from our motif analysis (**Table S2)**. After exclusions, we annotated the binding specificities of 57 unique lectins.

Binding motifs were annotated using a combination of two different approaches. First, we applied machine learning to identify the predominant binding motif(s) for each lectin. As input features for machine learning analysis, we curated a collection of 68 motifs (e.g., Blood Group B, terminal sialosides, etc.) seen in the CFGv5 (**Table S1)** and used these in tandem with all observed mono- and disaccharide motifs, including all observed linkages. These were used to generate binding rules and associated p-values (**Tables S3 & S4**) that best explained the experimental results. In all our analyses, the rules observed with machine learning were considered the predominant binding rules.

The machine learning analysis only gives a part of the picture, as only previously specified features can be detected as relevant. To complement our machine learning approach, we generated a combined Z-score analysis of the glycan microarray data and used that for further manual annotation (**Table S5**). Z-scores measure the deviation of individual glycans as binders (as reflected in fluorescence) from the mean. Using this metric as a measurement of binding assumes that the majority of glycans on the array are not bound by the lectin and thus the mean fluorescence indicates no true binding. We used Stouffer’s Z-score method to combine datasets of multiple concentrations for each lectin tested. This gave a single metric (Zs) for lectin binding to each glycan [18]. We set Zs=1.645 as our threshold for binding as this corresponds to a one-tailed *p*-value = 0.05, i.e., the 95% confidence interval [19]. In evaluating a binding motif, we first applied the machine learning rules. We then examined glycans following the rules that either bound or did not bind (based on Zs-score) and looked for features that could account for the difference. We used this information to annotate the predominant binding specificity. We next looked at glycans that did not follow the machine learning rules, but were nonetheless bound based on Z-score analysis. We again looked for features that could account for binding and annotated these as additional binding motifs. Combining machine learning with manual annotation gave a more complete description of binding than either method alone.

### Overview of Glycan Binding Profiles

To analyze the potential overlap in glycan binding motifs of the commercially available lectins, we hierarchically clustered our Zs datasets via average linkage analysis using the Pearson correlation coefficient as our distance metric. The heatmap of the cluster is shown in **Figure 2**. From this heatmap we can clearly observe that lectins bind distinct subsets of the glycome, arguing against the idea that they lack specificity. Generally speaking, lectins clustered according to their major specificities.

**FIGURE 2.**
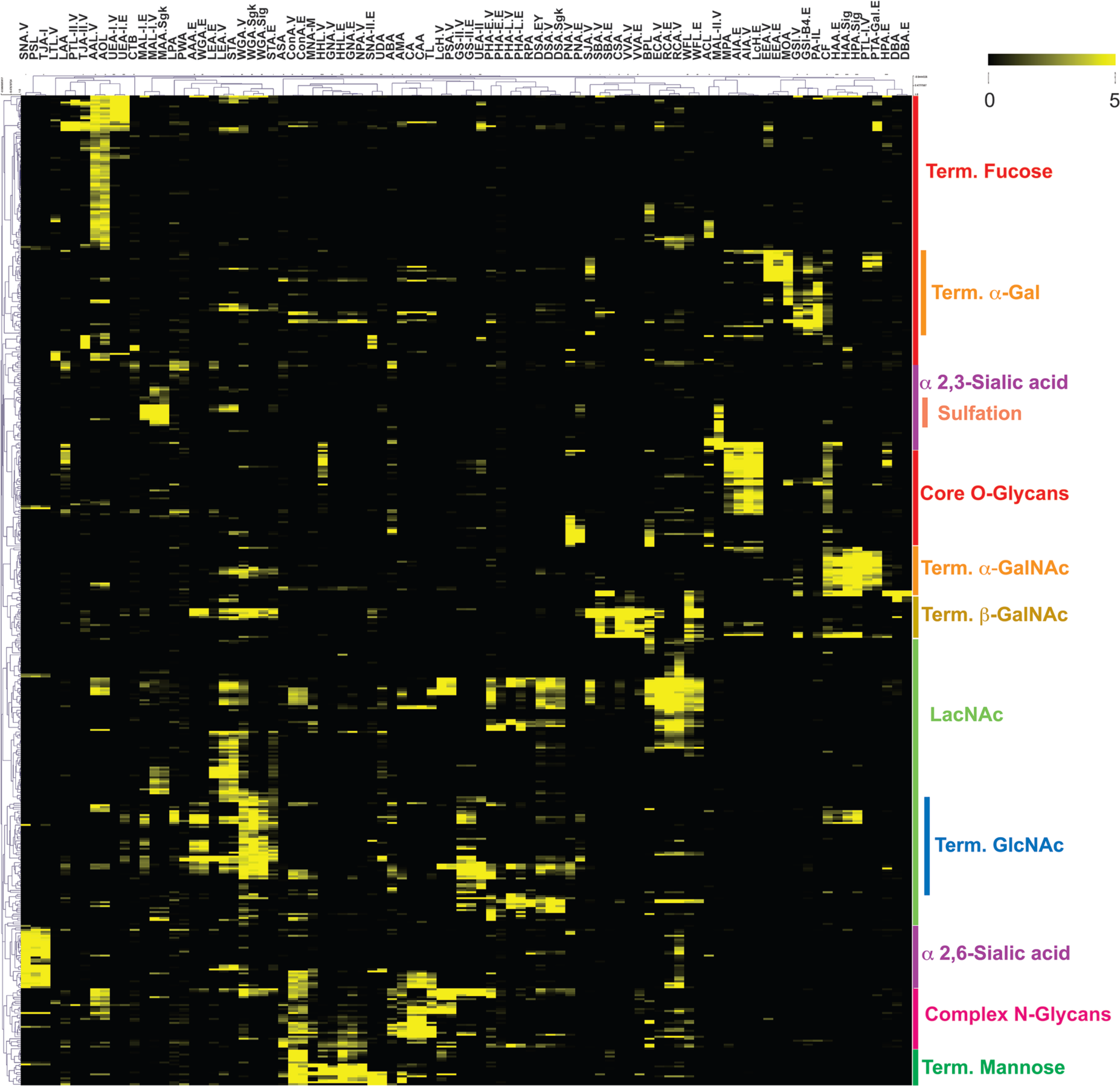
Heatmap of glycan binding data (Zs scores) for 90 lectins. Data was clustered using the Pearson correlation coefficient and average linkage analysis. Yellow indicates Zs > 1.645 (e.g. 95% confidence interval for binding), black indicates no significant binding. Rough annotations of glycan motifs are shown on the side. Term. = terminal.

In the following sections, we provide a detailed annotation of the glycan binding motifs of commercially available lectins revealed by our glycan microarray analysis. We have organized these by commonly referenced glycosylation motifs to make our analysis more useful to the scientific community. For each motif, we provide a table of lectins with the predominant and additional binding motifs outlined. We organized lectins using the following binding motifs: mannose (**Figure 3**), complex *N*-glycan (**Figure 4**), core *O*- glycans (**Figure 5**), fucose (**Figure 6**), sialic acid and sulfate (**Figure 7**), terminal *N-*acetylglucosamine (GlcNAc) and chitin (**Figure 8**), terminal galactose (Gal) and *N-* acetyllactosamine (LacNAc, **Figure 9**), and terminal *N-*acetylgalactosamine (GalNAc, **Figure 10**). In general, lectins are usually referred to by the Latin name of the plant from which they are derived, followed by the word agglutinin or lectin. This is typically shortened to a 3-4 letter acronym (e.g., wheat germ agglutinin = WGA). Where a lectin is commonly referred to as either the agglutinin or lectin, both acronyms are given. Detailed discussion of the binding specificities of the lectins can be found in the text below.

**FIGURE 3.**
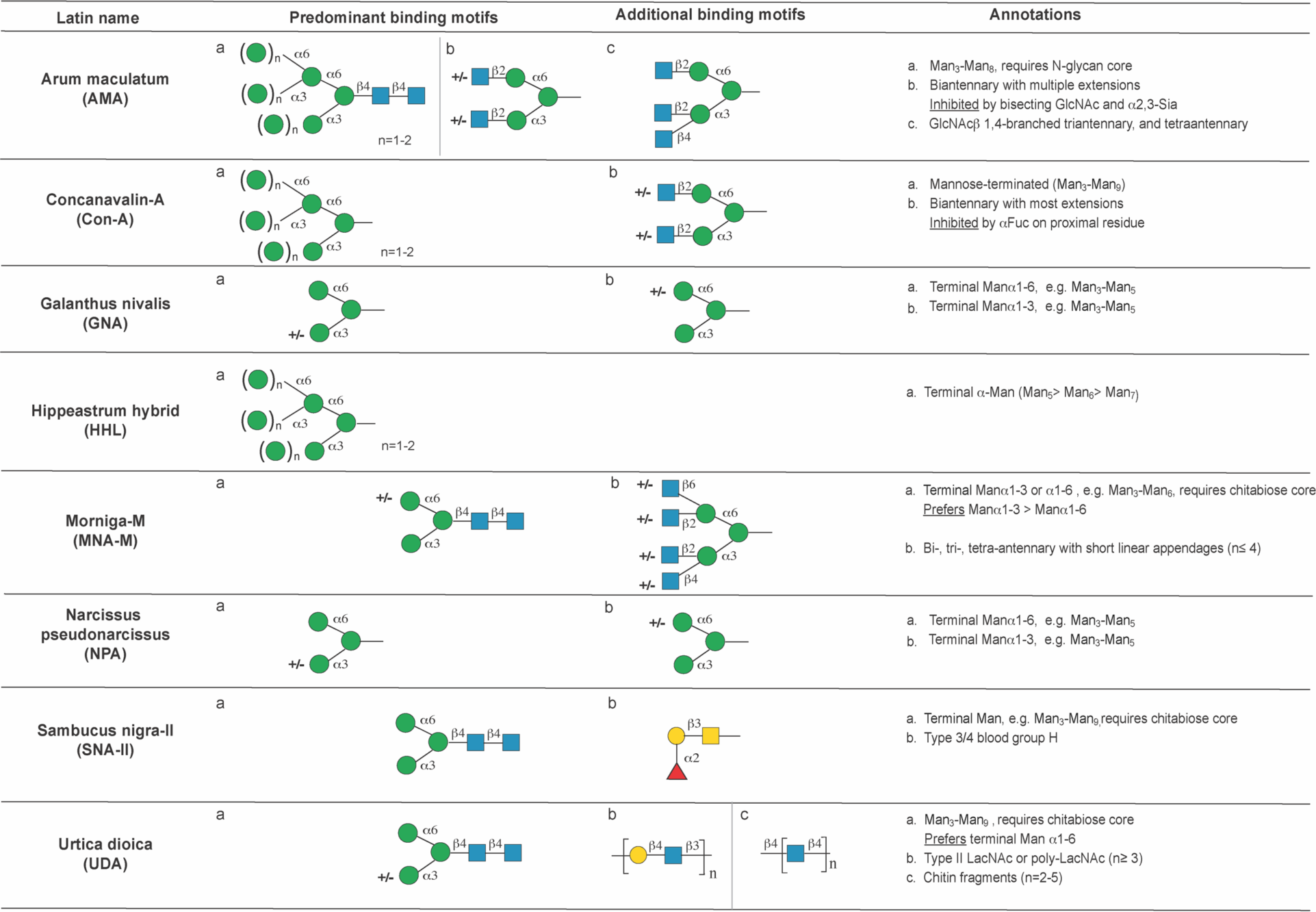
Binding motifs for mannose binding lectins. Abbreviations and Symbols: Mannose (Man, green circles), sialic acid (Sia, pink diamonds), *N-*acetylglucosamine (GlcNAc, blue squares), *N-* acetyllactosamine (LacNAc), fucose (Fuc, red triangles), galactose (Gal, yellow circles), *N-* acetylgalactosamine (GalNAc, yellow squares). The Symbolic Nomenclature for Glycans (SNFG) is used [1].

**FIGURE 4.**
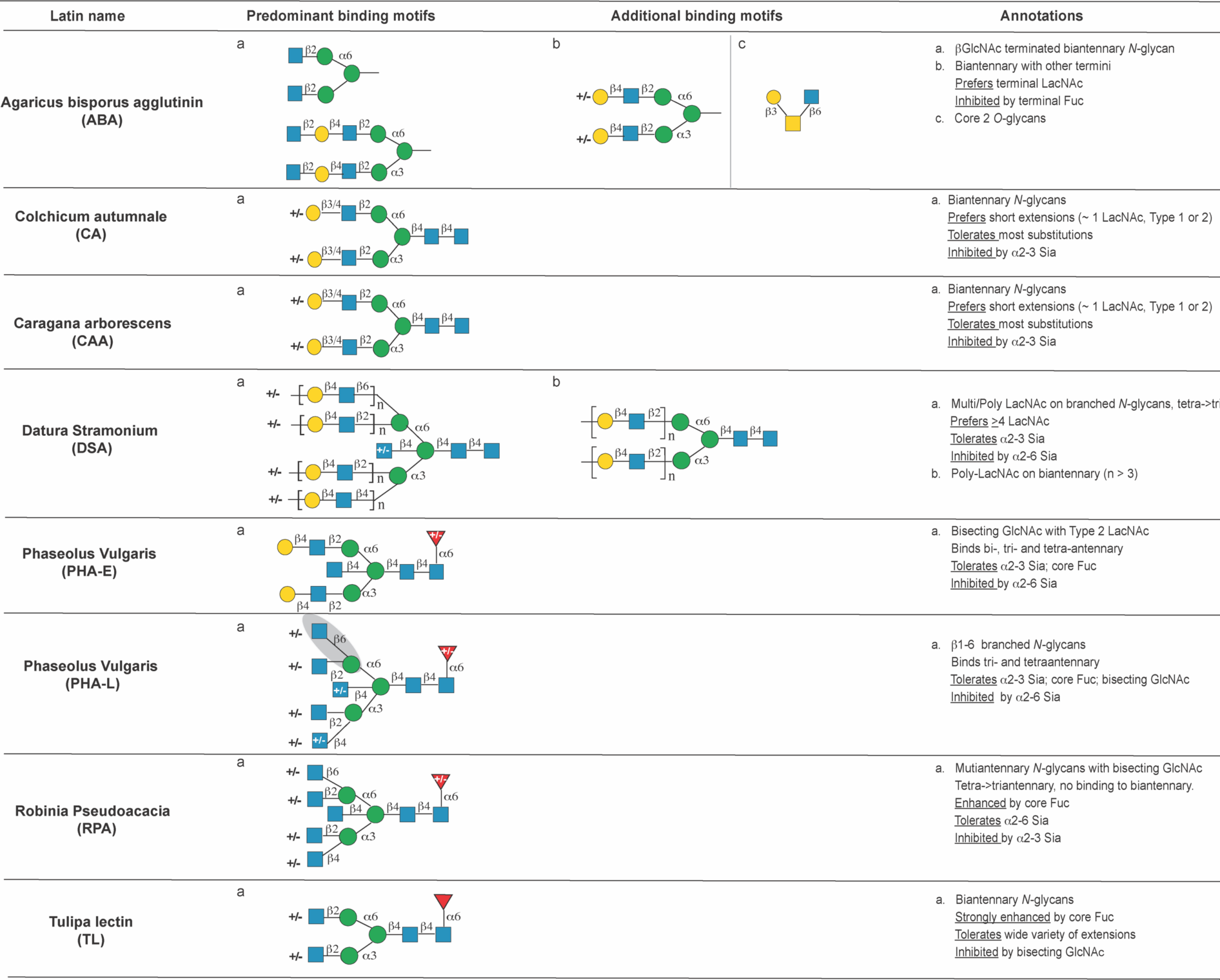
Annotation of lectins binding complex *N*-glycan motifs.

**FIGURE 5.**
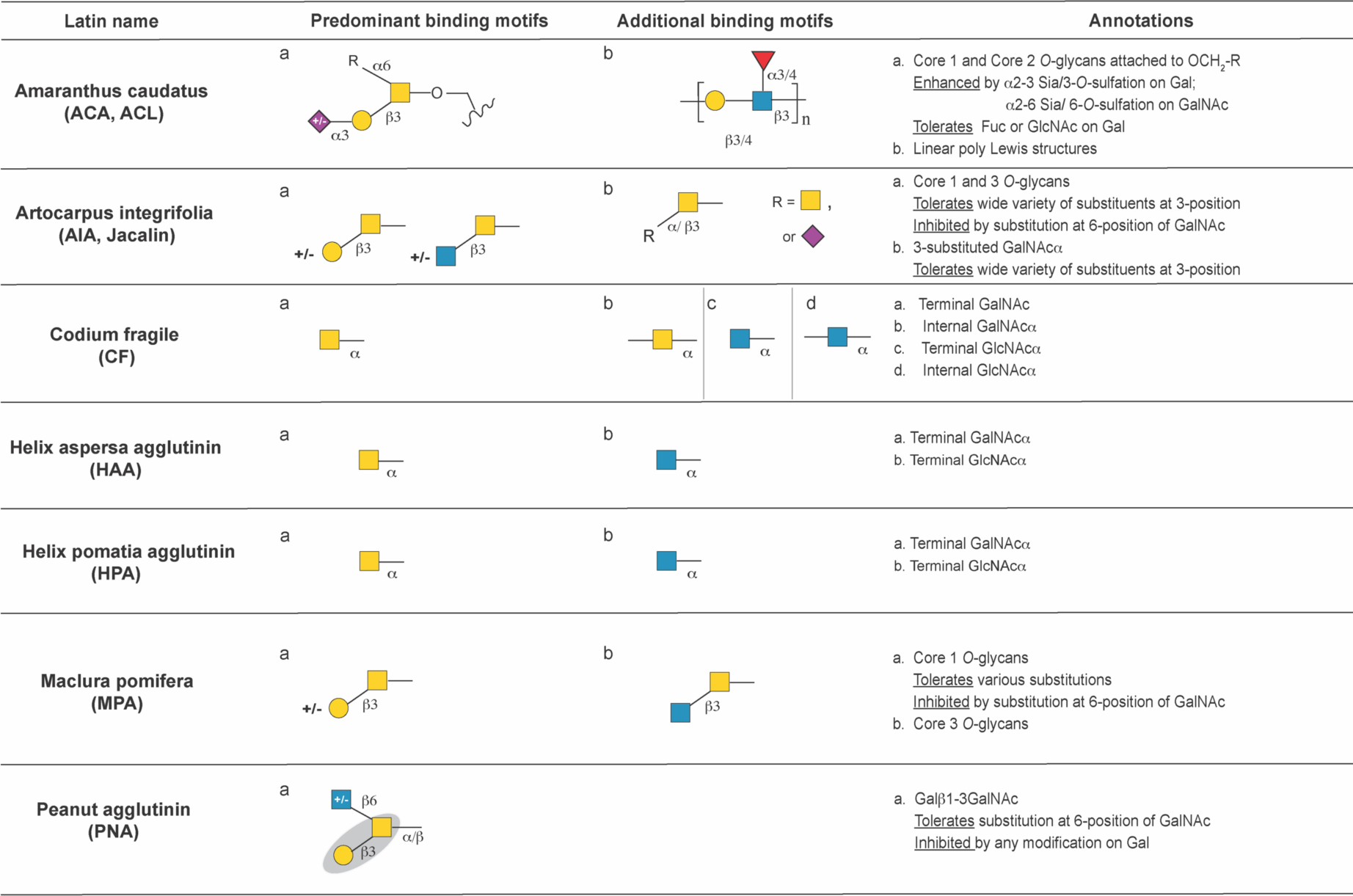
Annotation of lectins binding core O-glycan motifs.

**FIGURE 6.**
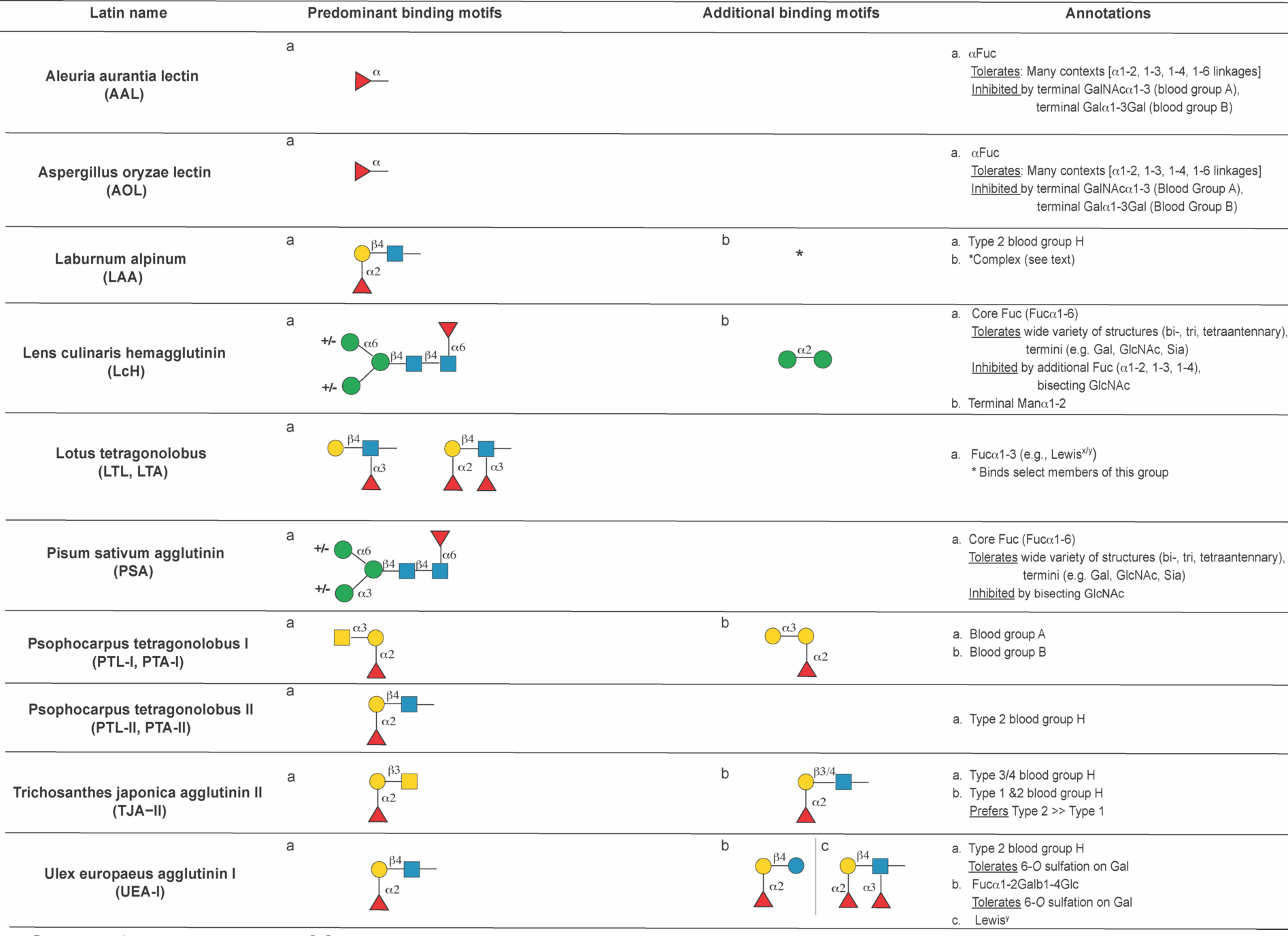
Annotation of fucose-binding lectins.

**FIGURE 7.**
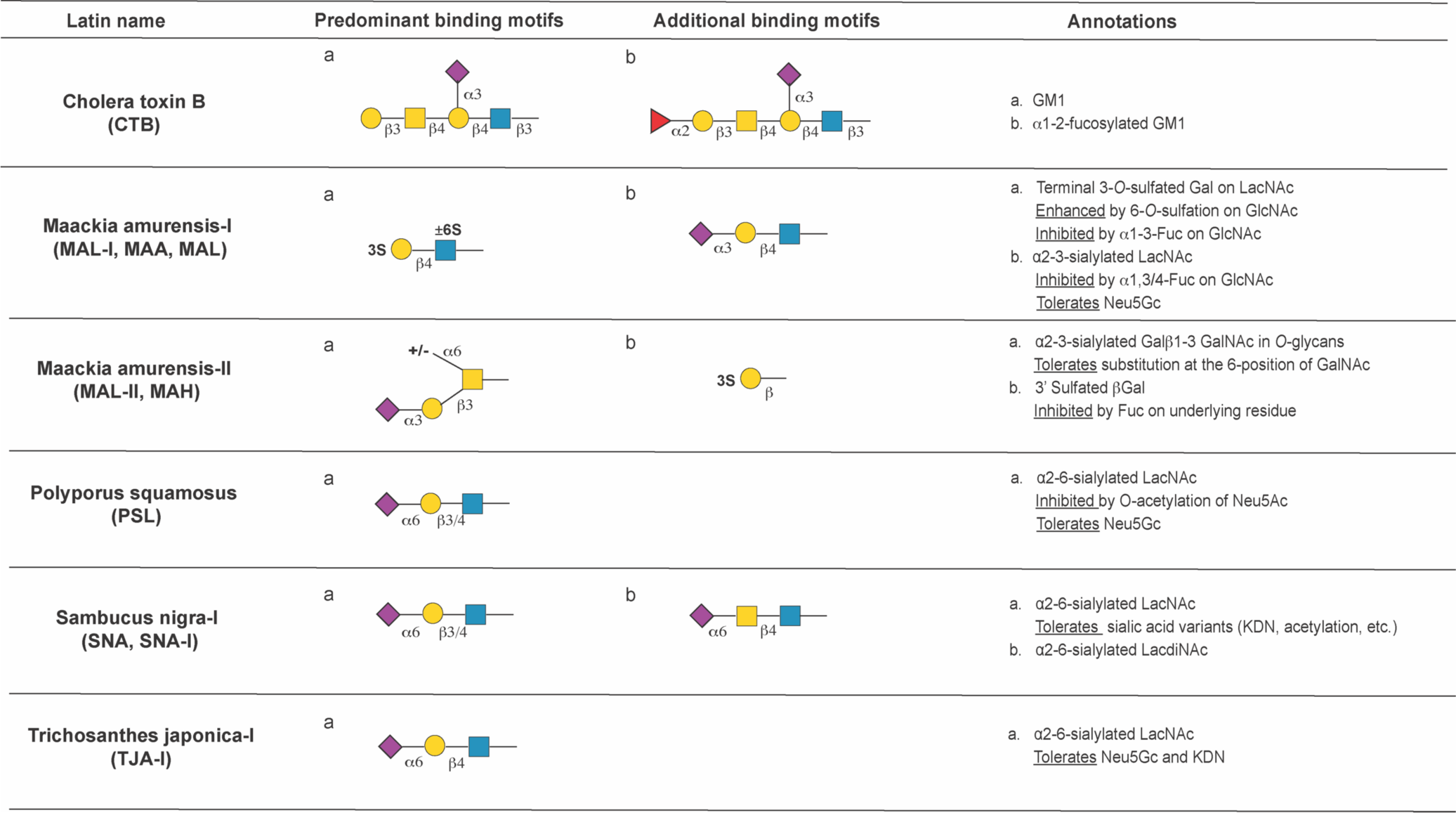
Annotation of sialic acid and sulfate binding lectins.

**FIGURE 8.**
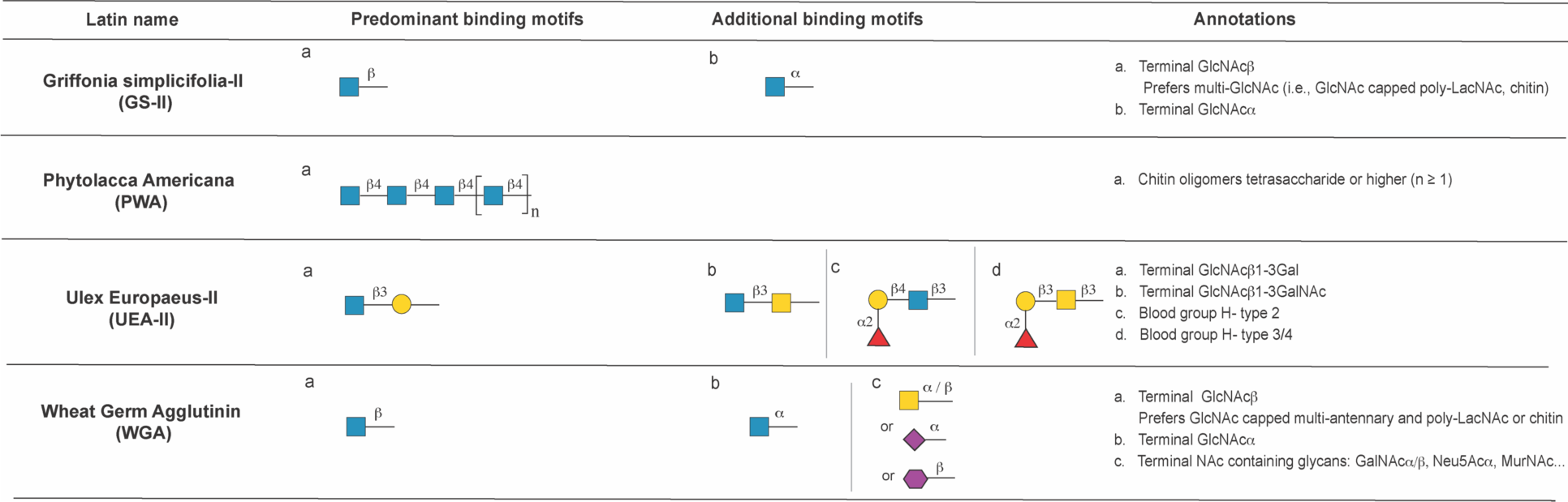
Annotation of GlcNAc and chitin binding lectins.

**FIGURE 9.**
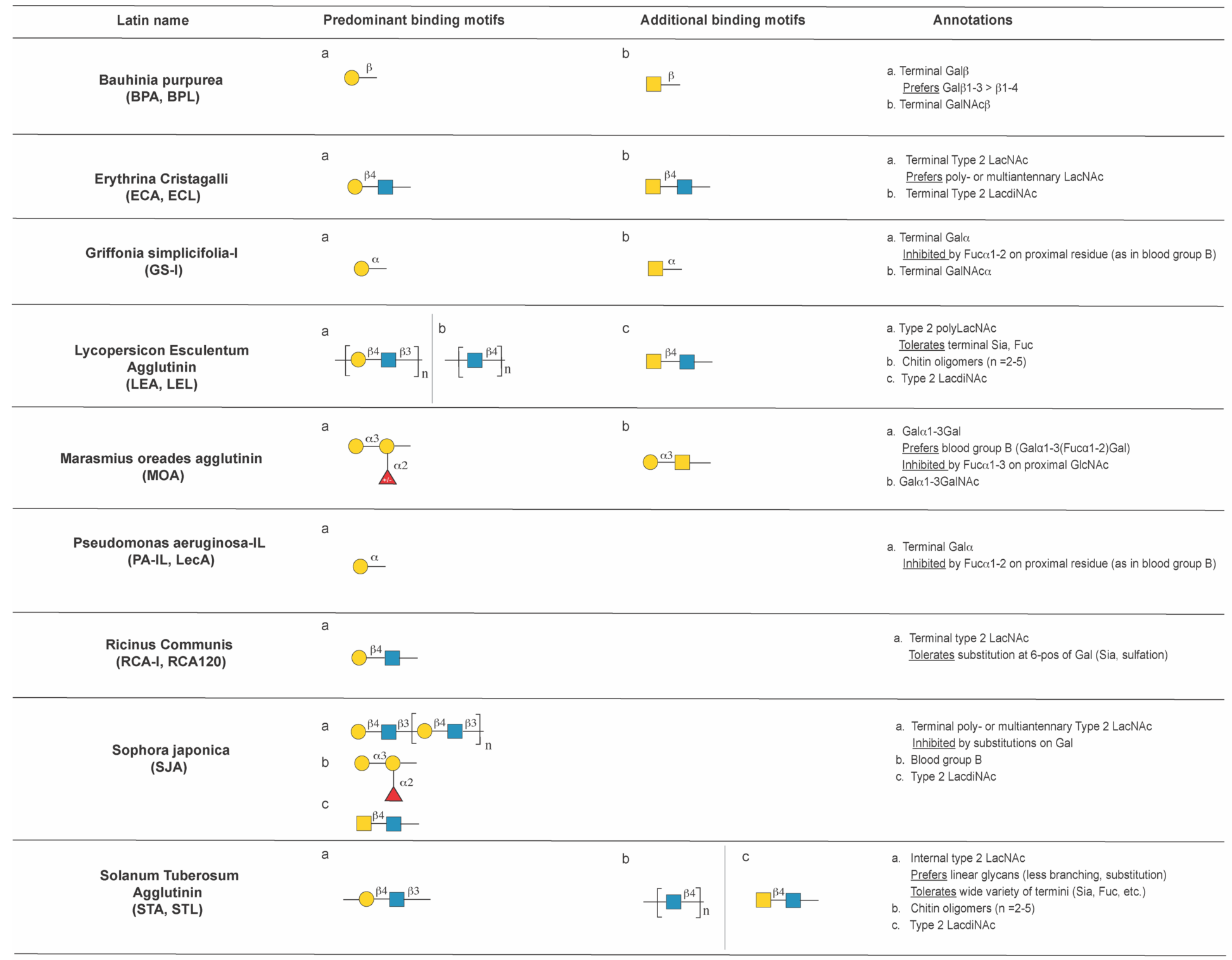
Annotation of Gal and LacNAc binding lectins.

**FIGURE 10.**
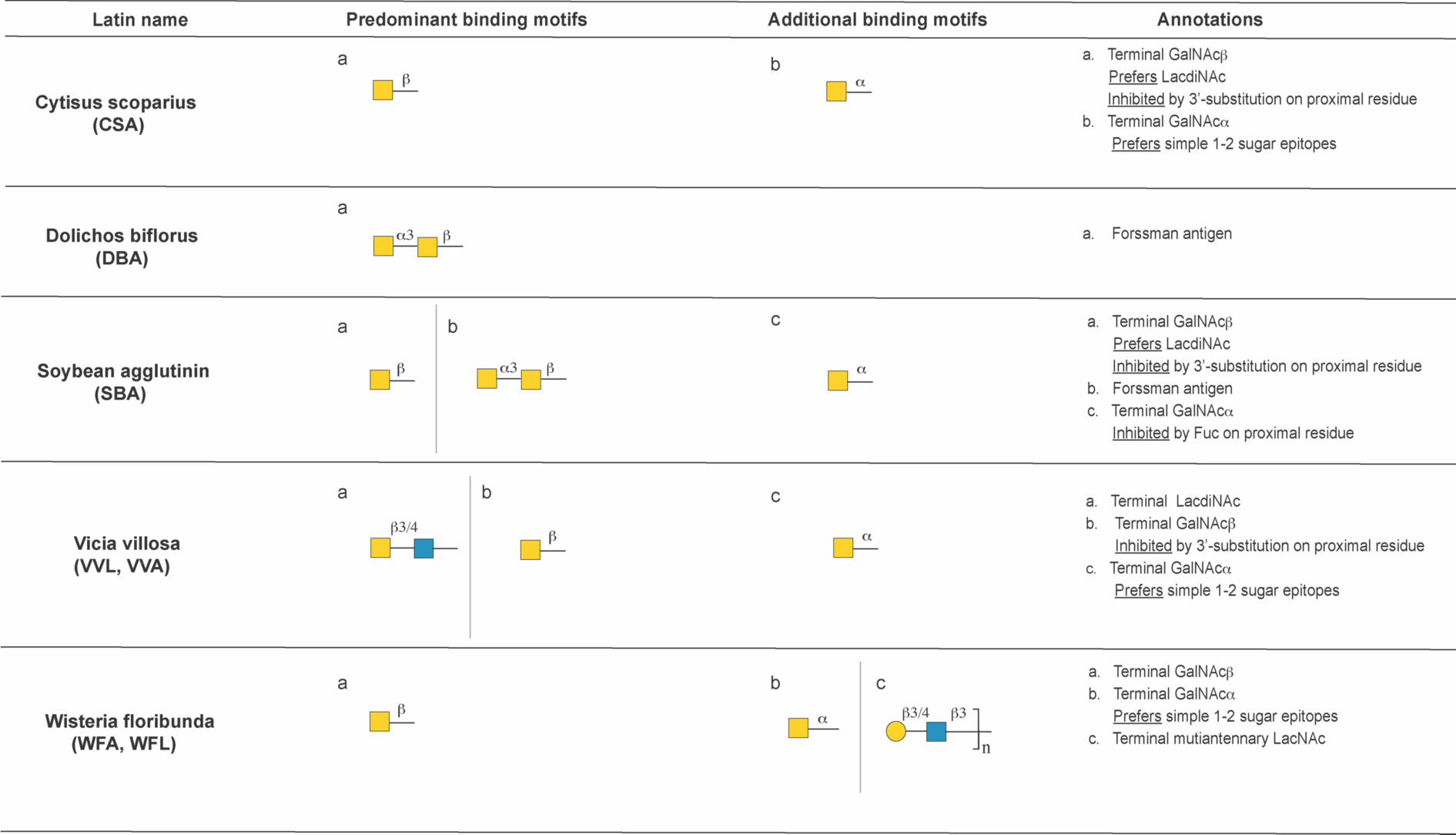
Annotation of GalNAc binding lectins.

### Mannose Binding Lectins

High mannose epitopes are among the least processed *N-*glycans, resulting from trimming of the Glc3Man9GlcNAc2-structure that is initially transferred co-translationally by oligosaccharyltransferase. Man7-Man9, i.e., high mannose, all contain terminal α1,2- mannose residues (**Figure S1A**). Further trimming of these structures results in oligomannose structures Man5-Man6, characterized by exposure of the trimannosyl core [Manα1-6(Manα1-3)Man]. This trimannosyl core is also exposed in hybrid *N-*glycan structures, although these are not represented on CFGv5. Mannose is also found as a direct modification of serines and threonines in non-canonical *O*-linked glycans (*O*- mannosylation) [20]. The following lectins predominantly recognized mannose-based epitopes on the array: *Arum maculatum* agglutinin (AMA)*, Concanavalin-A* (ConA), *Galanthus Nivalis* lectin (GNA, GNL)*, Hippeastrum Hybrid* lectin (HHA, HHL), *Morniga M* agglutinin (MNA-M), *Narcissus Pseudonarcissus* lectin (NPA)*, Sambucus Nigra* agglutinin II (SNA-II), *Urtica Dioica* lectin (UDA, **Figure 3**). A more detailed analysis of these lectins is given below.

### Arum maculatum (AMA)

The lectin from *Arum maculatum* (AMA) was isolated in 1995 on glycoprotein columns and noted to bind both *N-*acetyllactosamine (LacNAc) and mannose [21, 22]. In concordance with the original observations on this lectin, we observe two predominant glycan binding motifs: mannose-terminated *N-*glycans and biantennary structures (**Figure 3**). Perhaps due to the complexity of the two motifs no binding rules were identified by machine learning, but these motifs were clear in manual annotation. For the mannose-terminated glycans, AMA recognizes Man3-Man8. Recognition requires the chitobiose core (GlcNAcb1-4GlcNAc) and does not occur for mannose fragments. An exposed α1,3- or α1,6-mannosyl residue is observed in all binders. Man9, which does not fit these criteria, is not recognized. Biantennary glycans were also among the top binders. AMA can recognize triantennary *N-*glycans with β1,4-branching, and some tetraantennary with lower affinity. Although AMA tolerates a wide range of terminal structures (α2,6-sialic acid, fucose, Gal, GlcNAc, etc.) on bi-, β1,4-tri-, and tetra-antennary, its binding is inhibited in the presence of bisecting GlcNAc, or α2,3-sialic acid.

### Concanavalin-A (ConA)

First isolated in 1936, ConA, the hemagglutinin from jack bean, is perhaps the most commonly used lectin [23]. ConA is often annotated as a high-mannose binding lectin [24], although its true specificity is more complex. ConA from two sources (E.Y. Laboratories (EY) and Vector Laboratories (Vector)) were evaluated and had very similar binding patterns. Both vendors show terminal α-mannose as the predominant binding motif by machine learning (**Figure 3**). On the CFGv5 array, ConA recognizes Man3-Man9, along with truncated fragments, indicating that the core chitobiose is not necessary for recognition. Beyond mannose, ConA also recognized a wide range of biantennary *N-*glycans, tolerating multiple extensions (sialylated LacNAc, fucose, Gal, GlcNAc, etc.). However, binding is inhibited by α1,2- or α1,3-linked fucose attached to the proximal glycan at the termini (e.g., as in Galβ1,4 (Fuc α1,3) GlcNAc).

### Galanthus Nivalis lectin (GNA, GNL), and Narcissus Pseudonarcissus lectin (NPA, NPL)

GNA and NPA are both isolated from bulbs (snowdrop [25], and daffodil [26], respectively). On the CFGv5 array, the binding specificities of these two lectins overlap very closely and show no clear differences in binding determinants. Literature reports that GNA prefers terminal α1,3-linked [27], whereas NPA prefers α1,6-linked mannose [28]. machine learning identifies terminal Manα1-6 as the top motif for recognition by both lectins (**Figure 3**). Terminal Manα1-3 is also recognized, however, it exhibits weaker binding. The preference for exposed Manα1-6 or Manα1-3 argues that these lectins may recognize hybrid *N*-glycans, which are not present on this array. Man3-Man5 are preferred over Man6-Man8, which bear fewer available terminal α1,3- or α1,6- mannoses. Little (GNA) or no (NPA) binding is observed to terminal α1,2-linked mannose on the CFGv5 array. Our analysis of these lectins indicates short LacNAc terminated N-glycan epitopes are an additional binding determinant. GNA from two different sources showed highly similar binding patterns.

### Hippeastrum Hybrid lectin (HHA, HHL)

Isolated from amaryllis bulbs, *Hippeastrum hybrid* lectin (HHA, HHL) is known to bind terminal mannosyl residues or oligosaccharides containing either α1,3- or α1,6-linked mannosyl units [26]. Consistent with this, machine learning shows that HHL from two sources (EY and Vector) predominantly bind terminal *α*-mannose (**Figure 3**). This lectin shows the highest affinity for the mannose core trisaccharide (Manα1,6(Manα1,3)Man) and does not require the chitobiose termini. Man5-Man8 are bound, with Man5>Man6>Man7>Man8. Minor recognition of a smattering of LacNAc terminated structures is also observed, however, bound structures differ between the two preparations. HHL from Vector, but not EY, also recognizes several *O*-glycan structures (core 1 and core 3).

### Morniga-M (MNA-M)

One of two lectins isolated from the bark of the black mulberry tree (*Morus nigra*), Morniga M (MNA-M) was identified as a mannose specific lectin. Further characterization by frontal affinity chromatography found it to prefer Man3-Man6, with unsubstituted Manα1,3Man in the best binders [29]. Our analysis of this lectin is in keeping with this earlier work. The best binders for MNA-M all contain an unsubstituted Manα1-3 or Manα1-1. Machine learning shows a requirement for the chitobiose core (**Figure 3**). Binding to mannose fragments is not observed. Terminal Manα1-3 is preferred. We observed additional binding to bi-, tri- and tetraantennary *N-*glycans with short linear appendages (n=4 or less). It has been previously reported that core fucosylation inhibits its binding [30], however, this is not observed on the array.

### Sambucus Nigra Agglutinin II (SNA-II)

SNA-II is one of several lectins to be isolated from elderberry bark [31]. SNA-II is reported to be a terminal GalNAc/Gal binder [32], however these studies used only a small number of simple sugar structures (mono- and disaccharides) to probe the binding motif. Our machine learning analysis indicates that the top motif for SNA-II is terminal mannose (Man3-Man9) containing the chitobiose core as an essential component (**Figure 3**). Unlike ConA, this lectin does not recognize mannose fragments. This lectin also binds Type 3 & 4 blood group H antigens (e.g., Fucα1,2Galβ1,3GalNAc). Some binding is also observed to terminal Type 2 LacdiNAc (GalNAcb1,4GlcNAc).

### Urtica Dioica (UDA)

UDA, derived from stinging nettle, is annotated as a GlcNAc binding lectin [33].

Machine learning reveals that its predominant binding is terminal Manα1,6 (**Figure 3**). However, UDA will recognize Man3-Man9 with the chitobiose core. Like SNA-II, it will not bind mannose fragments. Chitin fragments (i.e., (-GlcNAcβ1,4GlcNAc-)n) and polymers of Type 2 LacNAc (n≥3 disaccharides) are also bound, but at a lower apparent affinity. In keeping with other work, our data points to a dual specificity for this lectin, with recognition sites for both mannose and LacNAc/chitin [34].

### Complex N-Glycan Binding Lectins

In *N-*glycan processing, high mannose *N-*glycans are trimmed to Man5 and a β1,2- GlcNAc is appended to the exposed core α1,3-mannose by MGAT1, forming hybrid structures (**Figure S1**). The trimannosyl epitope on the core α1,6-mannose is removed by MAN2A and the core α1,6-mannose branch further elaborated by MGAT2 to give biantennary *N-*glycans. ConA and AMA recognize biantennary *N-*glycans, however, as it is not their predominant binding motif, they are discussed with the high mannose lectins (**Figure 3**). Lectins from *Agaricus bisporus* (ABA, ABL)*, Colchicum autumnale* (CA), *Caragana arborescens* (CAA) and *Tulipa* lectin (TL) predominantly recognized biantennary *N-*glycans epitopes on the array (**Figure 4**)

Biantennary structures can be further elaborated to tri-and tetraantennary structures. In triantennary glycans, branching can occur at either the β1,4-position (GlcNAcβ1,4Manα1,3Man), controlled by the enzymes MGAT4A/B, or the β1,6- position (GlcNAcβ1,6Manα1,6Man), controlled by MGAT5/B, on the trimannosyl core [35]. Additionally, a GlcNAc can be added β1,4 to the central mannose of the core resulting in the bisecting GlcNAc motif and fucosylation of the chitobiose core is common (**Figure S1**). Several lectins are specific for these complex epitopes including: *Datura Stramonium* (DSA), *Phaseolus Vulgaris-*E (PHA-E), *Phaseolus Vulgaris*-L (PHA-L), *Robinia Pseudoacacia* (RPA). Core fucose binding lectins are included in **Figure 6**. A more detailed analysis of the lectins binding complex *N-*glycan epitopes is given below (**Figure 4**).

### Agaricus bisporus agglutinin (ABA, ABL)

Agaricus bisporus agglutinin (ABA, ABL), isolated from white button mushrooms, is thought to bind predominantly to *O*-glycans [36]. In contrast, our machine learning analysis identifies agalactosylated (GlcNAcβ-terminated) biantennary *N*-glycans as its predominant binding motif (**Figure 4**). In keeping with this, work by Hirabayashi and coworkers discovered that ABA displays dual-specificity, binding both agalactosylated biantennary and *O*-glycans [37]. ABA also recognizes biantennary *N*-glycans with other termini, especially terminal LacNAc, however, this binding is inhibited by fucosylation on or near the termini. Although literature indicates ABA binds to Core 1 [37], our analysis shows that Core 2 is preferred. We saw little binding to Core 1 epitopes on the array. Overall, our analysis is consistent with the dual binding specificity of ABA [37].

### Colchicum autumnale (CA) and Caragana arborescens (CAA)

CA, from meadow saffron, and CAA, the major lectin from the pea tree *Caragana arborescens* are both annotated as GalNAc binders [38–40]. These two lectins show almost identical binding patterns by machine learning (**Figure 4**). Both lectins bind bi- antennary *N*-glycans with short extensions (i.e., ∼1 LacNAc, either Type 1 or 2). CA and CAA tolerate most substitutions (e.g., fucose or α2,6-sialic acid), but binding is inhibited by α2,3-sialic acid. Although terminal blood group B is bound, blood group A is not recognized. We observe no recognition of GalNAc or Forssman type epitopes. Minor binding to tri- and tetra-antennary N-glycans with short chains is also observed.

### Datura Stramonium (DSA)

Isolated from Jimson weed (*Datura stramonium*), DSA was initially identified as a chitin-binding lectin [41], although further characterization revealed preferential binding to polylactosamine (polyLacNAc) and β1,6-branched *N-*glycans [42, 43]. Our analysis is largely consistent with these findings. Machine learning indicates that the predominant binding motifs are branching structures with four or more LacNAc repeats in total, either on different multiantennary branches, or polyLacNAc chains (n≥3 repeats) on biantennary *N-*glycans (**Figure 4**). Tetraantennary *N-*glycans containing Type 2 LacNAc are preferred to triantennary with similar epitopes, indicating that the binding affinity for DSA increases with higher branching. Both and β1,4- and and β1,6-triantennary structures are recognized. Biantennary *N-*glycans with long polyLacNAc chains (n≥3 LacNAc residues) are weaker binders. Bisecting GlcNAc and terminal α2,3-sialic acids are tolerated but α2,6 sialic acid inhibits lectin binding. DSA from three different sources (Vector, EY, Seikagaku) showed consistent binding patterns. It should be noted that on the CFG v5.0 array there is no representation of Type 1 LacNAc of a similar length or presentation. Thus, no conclusion could be made about whether DSA binds glycans containing Type 1 LacNAc residues.

### Phaseolus Vulgaris- erythroagglutinating (PHA-E) and Phaseolus Vulgaris- leukoagglutinating (PHA-L)

*Phaseolus Vulgaris*, also known as kidney bean, has at least five isolectins [3, 44, 45]. The two main isolectins that have been studied, PHA-E and PHA-L are so named due to their erythrocyte agglutinating (E) and leukocyte agglutinating (L) properties. Both lectins bind complex *N-*glycan epitopes with PHA-E annotated as a bisecting GlcNAc specific lectin [46] and PHA-L as a β1,6-branched *N-*glycan binder [47][46]. Our analysis is consistent with these annotations.

PHA-E from two sources (EY, Vector) show overlapping binding patterns. Both preparations of PHA-E have similar preferences for bisecting GlcNAc containing glycans (**Figure 4**). Core fucosylation and terminal α2,3-sialic acid structures are well tolerated, but α2,6-sialic acid inhibits binding. The only bisected Type 1 LacNAc ligand on the array, a biantennary *N-*glycan, is not recognized by PHA-E. However, an almost identical structure containing Type 2 LacNAc is among the best binders, arguing that this lectin may discriminate between Type 1 and Type 2 motifs. While both EY and Vector lectins showed similar preferences for bisecting GlcNAc, the preparation from Vector Laboratory recognizes a broader range of ligands including biantennary and triantennary structures without bisecting GlcNAc.

The machine learning analysis of PHA-L from EY and Vector shows that both preparations bind preferentially to β1,6-branched *N-*glycans, in keeping with the known binding specificity (**Figure 4**). Binding of β1,6-triantennary structures is inhibited by α2,6- but not α2,3-sialic acid. Bisecting GlcNAc and core fucose are tolerated. Although β1,6- branched glycans containing Type 2 LacNAc structures are well recognized, there is insufficient data to claim clearly whether or not Type 1 LacNAc is a ligand. Preparations from EY, but not Vector, showed additional binding to select bisected biantennary *N-* glycans. This may be due to differences in isolation techniques for this isolectin from natural sources, or contamination with PHA-E.

### Robinia Pseudoacacia (RPA)

*Robinia Pseudoacacia* or Black locust has two lectins that have been isolated from its seeds [48–50], in addition to several bark-derived lectins [3]. The commercially available preparation (E.Y. Laboratories) is a purified mixture of the seed proteins and is annotated as having complex specificity that is not inhibited by simple sugars. Machine learning shows that the principal binding determinant for RPA is multiantennary *N-*glycans containing a bisecting GlcNAc. Tetraantennary are preferred over triantennary structures and no binding is observed to biantennary structures. Core fucosylation enhances binding. Unlike other lectins in this group, binding is inhibited by α2,3-, but not α2,6-sialic acid residues.

### Tulipa lectin (TL)

The primary lectin isolated from the bulbs of tulips (*Tulip sp.,* TL) has not been well characterized. Initial experiments using agglutination assays with monosaccharides and glycoprotein-based inhibition studies showed complex sugar specificity [51]. Machine learning analysis indicates the principal binders for TL are biantennary *N-*glycans. The binding affinity is strongly enhanced by core fucosylation. This lectin is fairly permissive in the composition of extensions and termini, including sialic acid substituents, fucosylation, GlcNAc and LacNAc. However, bisecting GlcNAc has a negative impact on its binding affinity. TL shows lower affinity binding to triantennary *N*-glycans, but prefers β1,4-branched over β1,6-branched structures.

### Core O-glycan Binding Lectins

Unlike *N*-glycosylation, where a large oligosaccharide is transferred *en bloc* to asparagines on the nascent polypeptide, *O*-glycosylation begins with the transfer of a single sugar, *N-*acetylgalactosamine (GalNAc, **Figure S1**), onto serine or threonine. This glycan epitope, also known as the Tn antigen, can then be elaborated by a host of glycosyltransferases to make a variety of core structures. *O*-glycosylation is perhaps best studied on mucins, glycoproteins that have clustered *O*-glycan sites. These proteins contribute to a wide variety of biological systems, impacting everything from lung function to cancer progression [52]. Lectins that recognize *O-*glycans specifically include *Amaranthus Caudatus* (ACA, ACL), Peanut agglutinin (PNA), *Artocarpus integrifolia* (AIA, Jacalin), *Codium Fragile (CF), Maclura Pomifera* (MPA, MPL), Helix pomatia agglutinin (HPA) and Helix aspersa agglutinin (HAA) (**Figure 5**). *Maackia Amuerensis-II* (MAL-II), which binds sialic acid on *O-*glycans, is covered in **Figure 7**. A detailed analysis of these lectins is given below.

### Amaranthus Caudatus (ACA, ACL)

Isolated from the seeds of Tassel flower (*Amaranthus caudatus*), ACL is widely considered to bind T-antigen (Galβ1,3GalNAc-Ser/Thr) [53]. Our machine learning analysis shows that the best binders for ACL are core 1 and core 2 *O*-glycans, both of which contain Galβ1,3GalNAc (**Figure 5**). ACL shows a strong linker dependency in its recognition. No binding is observed when threonine (Sp14) is used as the linker. Instead, the unbranched propyl amino-linker Sp8 (-CH2CH2CH2NH2) is strongly preferred. This is suggestive that ACL binds serine rather than threonine-linked *O-*glycans. However, as serine is not used as a linker on this array, it is not conclusive. Binding is enhanced by sulfation or sialylation at the 3-position of Gal and/or at the 6-position of the core GalNAc. Binding also tolerates other substitution on Gal, including fucose. ACL also shows lower binding affinity to repeated Lewis structures (poly Lewis-x, n ≥ 2), or combinations of Le^x^, Le^y^ and Le^a^ antigens.

### Artocarpus integrifolia (AIA, Jacalin)

The lectin isolated from *Artocarpus integrifolia* (aka Jackfruit: AIA, Jacalin) [54] is widely considered a T-antigen binder. In keeping with this, machine learning reveals this lectin predominantly binds 3-substituted GalNAcα (e.g., Core 1 and Core 3 *O*-glycans, **Figure 5**). A wide variety of substituents are tolerated at the 3-position of GalNAc via either an α or β-linkage, including GalNAc, GlcNAc, Gal and longer oligosaccharides. However, binding is inhibited by any substitution at the 6-position of the core GalNAcα, precluding recognition of Core 2, 4, 6 and 7-*O*-glycans. AIA from EY and Vector display identical binding patterns.

### Codium fragile (green marine algae, CF)

Isolated from the green marine algae, *Codium fragile* (CF) is reported to be a GalNAcα specific lectin [55]. Machine learning analysis is consistent with these reports. The predominant binder is terminal GalNAcα and it binds all epitopes containing this glycan including Tn and Blood Group A (**Figure 5**). CF also recognizes internal GalNAcα structures including Core 1 and 3 *O*-glycans. However, unlike AIA, this lectin is insensitive to substitutions of the core GalNAc at the 6-position, as seen in core 2 and core 4 *O*- glycans, and the sialyl-Tn antigen. The lectin will also bind glycans containing terminal and internal GlcNAcα, which are not seen in mammals.

### Helix pomatia agglutinin (HPA) and Helix aspersa agglutinin (HAA)

*Helix pomatia* agglutinin (HPA), found in the albumen gland of the roman snail [56], and an agglutinin from garden snail *Helix aspersa* (HAA) [57], are both commonly used as probes for αGalNAc (Tn antigen), an antigen with strong associations to cancer [58].

The two lectins exhibit almost identical binding patterns. Machine learning shows a strong preference for GalNAcα-terminated oligosaccharides (**Figure 5**). Both lectins bind terminal GalNAcα on Tn antigen and Blood Group A. Binding to GlcNAcα is also observed, arguing that the stereochemistry at the 4-position of the terminal sugar is not essential to binding. HAA and HPA from two sources (EY, Sigma) show similar specificities across all four preparations.

### Maclura Pomifera (MPA, MPL)

The lectin from *Macluria pomifera* (osage orange: MPA) [59] is considered a T-antigen binder. Machine learning shows that this lectin predominantly binds Core 1 *O*- glycans (**Figure 5**). Similar to AIA, it tolerates a wide variety of Core 1 *O*-glycans but binding is inhibited by substitution at the 6-position of the core GalNAcα (e.g., Core 2). Binding is also observed to Core 3 epitopes.

### Peanut agglutinin (PNA)

Isolated from the peanut (*Arachis hypogaea)*, PNA is commonly considered a T-antigen (Galβ1,3GalNAc) binding lectin [60]. Machine learning shows that terminal Galβ1,3GalNAc is indeed the preferred ligand, although the GalNAc residue can have either α- or β-linkages (**Figure 5**). This lectin is insensitive to substitutions of the core GalNAc at the 6-position, as is seen in various core 2 *O*-glycans. However, binding is inhibited by any substitution on the Gal termini, arguing the requirement for unhindered access to terminal Galb1-3GalNAc for its binding. PNA from Vector and EY have overlap, but their binding patterns are not identical. In the EY preparation overall fewer binders and low levels of binding to chitin epitopes are observed. In contrast, binding to GlcNAc terminated LacNAc is observed in the Vector preparation.

### Lectins binding structures common to N- and O-glycans

*N*- and *O*-glycans carry a variety of epitopes beyond their core structures. These include terminal epitopes such as sialic acid and internal ones such as polyLacNAc. These glycan modifications confer unique functional properties to oligosaccharides and impact a variety of biological processes from immune recognition to neurogenesis. In this section we discuss lectins that bind a range of epitopes found on *N-* and *O-*glycans. These include fucose (**Figure 6**), sialic acid and sulfate (**Figure 7**), terminal GlcNAc and chitin (**Figure 8)**, terminal galactose and LacNAc (**Figure 9**), and terminal GalNAc (**Figure 10**).

### Fucose binding lectins

Often considered a terminal modification, fucose is observed in diverse structural contexts within glycans and impacts an array of biological functions. Core fucosylation, which in mammals is exclusively α1,6-linked to the asparagine linked GlcNAc in *N-* glycans, is a commonly observed modification. This modification requires trimming and elongation of the *N-*glycan to hybrid structures and is observed on both hybrid and complex *N*-glycans (**Figure S1**) [20]. Core fucosylation of antibodies inhibits antibody-dependent cell-mediated cytotoxicity [61]. α1,3- and α1,4-fucosylation are most commonly studied in the context of Lewis structures, including sialyl-Lewis^x^, which plays a role in inflammation [62]. α1,2-fucose is a key component of blood group antigens. Lectins that predominantly recognize fucosylated glycans include *Aleuria aurantia* lectin (AAL), *Aspergillus oryzae* lectin (AOL), *Laburnum alpinum* lectin (LAA), *Lens culinaris* hemagglutinin (LcH, LcA), *Lotus tetragonolobus* lectin (LTL), *Pisum sativum* agglutinin (PSA), *Psophocarpus tetragonolobus* lectin-I (PTL/PTA I) and -II (PTL/PTA II), *Trichosanthes japonica* agglutinin II (TJA−II), and *Ulex europaeus* agglutinin-I (UEA-I) as shown in **Figure 6**.

### Aleuria aurantia lectin (AAL) and Aspergillus oryzae lectin (AOL)

The fungal lectins from *Aleuria aurantia* (AAL) [63] and *Aspergillus oryzae* (AOL) [64] are both known to bind fucose in many forms. Machine learning analysis shows that both lectins primarily bind to *α*-linked fucose and have very similar specificities. They can bind fucose residues linked to oligosaccharides at the 2-, 3-, 4-, or 6-position (**Figure 6**). However, machine learning also revealed unexpected subtleties in their binding. Neither lectin recognized α1,2-fucose in the context of full epitopes of blood group A [GalNAcα1,3(Fucα1,2)Galβ1,3/4GlcNAc, BGA] or blood group B [Galα1,3(Fucα1,2)Galβ1,3/4GlcNAc, BGB]. There are also subtle differences between the two lectins. AAL shows a preference for Fucα1,2-terminated structures on Type 2 over Type 1 LacNAc, a finding not observed in AOL. In addition, only AOL was able to recognize Type 3/4 blood group H antigens (Fucα1,2Galβ1,3GalNAc).

### Laburnum alpinum Lectin (LAA)

*Laburnum alpinum* lectin (LAA) comes from the Scotch laburnum tree and is known as a blood group H binder [65]. Machine learning shows it prefers the Type 2 blood group H epitope on *N*-linked glycans (**Figure 6**). It does not bind this epitope when presented on an *O*-glycan core. LAA is also reported to recognize N-acetylglucosamine/chitobiose [66]. In keeping with this, we observed weaker binding to a complex mixture of other glycans. The majority of these structures contained a combination of β1,3-linkages and N-acetyl groups (e.g., GlcNAc β1,3Gal (in polyLacNAc), Type 1 LacNAc, etc.).

### Lens culinaris hemagglutinin (LcH, LcA)

*Lens culinaris hemagglutinin* (LcH, LcA) from lentils [67] is reported to be a core fucose specific lectin. Machine learning confirms that core fucose (α1,6-Fuc) is the major binding determinant (**Figure 6**). However, the presence of additional fucosylation in the glycan (e.g,, α1,2-, α1,3-, α1,4-Fuc) or bisecting GlcNAc inhibits binding. The lectin can tolerate a wide variety of terminal epitopes (e.g., α2,6- and α2,3-sialic acid, Gal, GlcNAc, and GalNAc), and a variety of branched structures (i.e., bi-, tri-). This is in contrast to previous work using frontal affinity chromatography that shows no binding for core fucosylated triantennary glycans [68]. In that work, only core fucosylated triantennary structures branched at the β1,4-, but not the β1,6-position, were examined [68]. The single β1,4- triantennary core fucosylated eptiope on CFGv5 does not bind LcH, however multiple examples of β1,6- triantennary core fucosylated epitopes are bound. Thus, it is likely that these lectins discriminate between β1,4- and β1,6-branching, although this cannot be definitively determined. LcH also binds mannose structures containing terminal Manα1,2, although these are weak binders, in line with previous reports [68].

### Lotus tetragonolobus lectin (LTL)

*Lotus tetragonolobus* lectin (LTL, Lotus) was originally isolated from lotus seeds as a fucose binding lectin [69]. Originally annotated as a fucose-binding anti-H(O) lectin [70], more recent analysis has identified Le^x^ as its main recognition motif [71]. No rule was identified by machine learning, however α1,3-fucose was significantly enriched in the subset of LTL binding glycans (**Figure 6, Table S4)**, in line with previous work. The α1,3- fucosylated glycans Le^x^ and Le^y^ were bound, as was 6’sulfo-SLe^x^. However, many α1,3- fucosylated GlcNAc containing glycans were not bound, including SLe^x^, and the rules governing binding of this subset were not clear. No binding was observed to glycans bearing only α1,2-fucosylated glycans.

### Pisum sativum agglutinin (PSA, PSL)

Isolated from peas, *Pisum sativum* agglutinin (PSA, PSL) was originally reported as a mannose binding lectin [72] [73], although it is currently thought to bind core fucose. Its binding is closely related to LcH. Machine learning confirms core fucose (α1,6-Fuc) is the major binding determinant and it tolerates a wide variety of structures (**Figure 6**). The presence of bisecting GlcNAc inhibits binding. Mannose binding was not observed on this array, but has been widely reported for this lectin [74].

### Psophocarpus tetragonolobus lectin-I (PTL-I/PTA-I) and -II (PTL-II/PTA-II)

*Psophocarpus tetragonolobus* lectin *-*I (PTL-I) and -II (PTL-II), isolated from winged bean seeds, are lectins with different carbohydrate binding specificities [75]. PTL-I is reported to be an *α*GalNAc-specific lectin [76]. Machine learning indicates that blood group A trisaccharides [GalNAca1,3(Fuca1,2)Gal] are the predominant binding motif for this lectin (**Figure 6**). The *α*GalNAc, however, is not strictly required as this lectin also recognizes blood group B, which has a terminal *α*Gal residue. Binding is somewhat inhibited by additional α1,3-fucosylation on internal GlcNAc residues.

In contrast to PTL-I, PTL-II is thought to specifically recognize blood group H [77]. In keeping with this, machine learning shows that PTL-II binds Type 2 blood group H epitopes, e.g., Fuca1,2Gal*β*1,4GlcNAc (**Figure 6**). It does not bind Le^y^.

### Trichosanthes japonica agglutinin II (TJA−II)

*Trichosanthes japonica* yields two distinct lectins, Trichosanthes japonica-I (TJA-I), a sialic acid binder covered in **Figure 7** and Trichosanthes japonica-II (TJA-II), which is thought to be specific for blood group H [78]. Machine learning identifies Type 3/4 blood group H, containing the Fucα1,2Galβ1,3GalNAc structure, as the predominant binding motif (**Figure 6**). Among the four types of blood group H antigens, H type 3/4 have the highest binding affinity, while Types 1 and 2 are weaker binders, with a preference order: Type 2 H >> Type 1 H.

### Ulex europaeus agglutinin-I (UEA-I)

The gorse plant, *Ulex europaeus* has two major lectins, *Ulex europaeus* agglutinin-I (UEA-I) and -II (UEA-II) [79][80]. Although these lectins are homologous, they show distinct binding specificities [81]. UEA-I, annotated below, is known as a fucose lectin (**Figure 6**), whereas UEA-II is considered a chitin-binding lectin (**Figure 8**).

UEA-I is well known to recognize the Fucα1-2Gal in fucosylated oligosaccharides [82]. In keeping with this, machine learning rules shows Type 2 blood group H (Fucα1,2Galβ1,4GlcNAc), as the predominant binding epitope (**Figure 6**). No binding is observed to Type 1 H epitopes (Fucα1,2Galβ1,3GlcNAc), indicating that the nature of the Gal linkage is important. The GlcNAc is less important, as epitopes containing Glc (e.g., Fucα1,2Galβ1,4Glc) or with substitutions on the GlcNAc (e.g., Le^y^) are tolerated. UEA-I tolerates sulfation on the 6-position of the terminal Gal.

### Sialic Acid and Sulfate Binding Lectins

Both sialic acid and sulfation bring a negative charge to glycans. Sialic acid (Sia), also known as *N*-acetylneuraminic acid (Neu5Ac), can be found in a variety of linkages (α2,3- , α2,6- and α2,8-) and is a terminal structure on *N-* and *O-*glycans and glycolipids. Abundant in the brain, sialosides modulate many processes, including neuronal migration, inflammation, and viral pathogenesis [20]. Sulfation is often found in glycosaminoglycans, such as heparin, but is also critical on *N-* and *O-* glycans and glycolipids. Sulfation has important roles in immunology, for example, 6’-sulfo SLe^x^ is a major ligand of L-selectin [83]. The following lectins bind to sialylated or sulfated structures on the array: *Cholera toxin B* (CTB), *Maackia Amurensis-I* (MAL-I, MAM, MAL), *Maackia Amurensis-II* (MAL-II, MAH), *Polyporus Squamosus* (PSL), *Trichosanthes Japonica-I* (TJA-I), and *Sambucus Nigra-I* (SNA-I) (**Figure 7**). The specificity of these lectins is discussed in detail below.

### Cholera toxin B subunit (CTB)

Although not a lectin, the B-subunit of Cholera toxin (CTB) is commonly used to stain for the ganglioside GM1 [Galβ1,3GalNAcβ1,4(NeuAcα2,3)Galβ1,4Glc] [84][85]. Due to both the complexity and specificity of the bound epitope, machine learning was unable to identify a significant binding rule. However, on CFGv5 only the glycosphingolipid epitopes, GM1 and fucosylated GM1 [Fucα1,2Galβ1,3GalNAcβ1,4(NeuAcα2,3)Galβ1,4Glc] were bound (**Figure 7**). This is in keeping with the known specificity of CTB.

### Maackia Amurensis-I (MAL-I, MAM, MAL) and -II (MAL-II, MAH)

Lectins isolated from the seeds of *Maackia amurensis* are commonly used to probe for α2,3 sialic acids [86]. Two lectins have been identified from *Maackia*, MAL-I (MAM, MAL, often referred to as MAA) and MAL-II (MAH). Although these lectins have similar amino acid sequences (86.2 % identity) [87], they have distinct binding specificities [86]. Machine learning analysis of MAL-I from three sources (E.Y. Laboratories, Vector Laboratories,and Seikagaku) shows that this lectin preferentially binds to terminal 3-O sulfated Gal on LacNAc (**Figure 7**). Although this has been observed in other studies [11, 86], it is contrary to the common usage of this lectin as a sialic acid binder. Binding is inhibited by fucosylation at the 3-position of GlcNAc (e.g., as in 3’*O*-sulfo Le^x^) and is enhanced by *O*-sulfation at the 6-position of GlcNAc. Although 3’*O*-sulfation is the strongest binding motif, only ∼30% of all binders are covered by this rule. In 2 of the 3 preparations of this lectin (Vector, Seikagaku), terminal *α*2,3-sialic acid on Type 2 LacNAc is observed as a second determinant. Binding is again inhibited by fucosylation at the 3, or 4-position of GlcNAc (i.e., MAA-I does not bind sialyl Le^x^ or sialyl Le^a^). A variety of sialic acid variants were tolerated, including Neu5Ac, 5-*N*-glycolyl neuraminic acid (Neu5Gc), 9O, 5N- diacetylated neuraminic acid (Neu5,9Ac) and KDN.

In contrast to MAL-I, the predominant binding determinant for MAL-II as defined by machine learning is α2,3-sialylated Galb1-3GalNAc in *O*-glycans (**Figure 7**). Various substitutions at the 6-position of GalNAc are tolerated, including sialylation (Neu5Ac or KDN), sulfation and GlcNAc, implying that the 6-position of GalNAc is not critical to recognition. This is in keeping with previous analysis of this lectin [86]. Similar to MAL-I, 3’O-sulfated Gal epitopes were observed to be an additional binding motif, however disulfation of galactose and/or fucosylation of the underlying residue are not tolerated.

### Polyporus Squamosus Lectin (PSL, note: this abbreviation is also used by Pisium Sativum Lectin)

Isolated from a mushroom, the main lectin from *Polyporus squamosus* is known to bind α2,6-sialylated LacNAc [88, 89]. Machine learning confirms this as the principal binding motif of PSL, and all glycans on the array containing this motif, regardless of context, are recognized (**Figure 7**). This includes the single example of α2,6-sialylated Type I LacNAc on the array. PSL tolerates Neu5Gc, but does not bind to O-acetylated sialic acid (Neu5,9Ac).

### Sambucus Nigra Agglutinin (SNA, SNA-I)

One of several lectins isolated from elderberry bark (*Sambucus nigra)* SNA-I, also known as SNA, is the most commonly used probe for α2,6-sialic acid [90]. SNA was thought to require a disaccharide of the structure Neu5Acα2,6Gal/GalNAc. Machine learning reveals α2,6-sialylated LacNAc as the predominant binding determinant (**Figure 7**). As with PSL, the single case of Type I α2,6-sialylation on the array is bound. α2,6- Sialylated LacdiNAc is also a binder. In contrast, no binding is observed to sialyl Tn antigen, in contravention of earlier literature [90]. A wide variety of α2,6-sialic acids are tolerated (e.g., Neu5Gc, KDN), in keeping with previous findings [91].

### Trichosanthes Japonica-I (TJA-I)

TJA-I, one of several lectins isolated from *Trichosanthes japonica*, is known to bind to Neu5Acα2,6Galβ1,4GlcNAc [92]. Machine learning confirms this specificity and identifies the predominant binding motif as α2,6-sialylated LacNAc (**Figure 7**). The strongest binders present this motif on multiple antennae. TJA-I tolerates Neu5Gc and KDN. Weak binding is observed to the α2,6-sialylated Type I LacNAc epitope.

### Terminal GlcNAc, and Chitin Binding Lectins

Terminal GlcNAc residues are common capping groups and are seen in a wide variety of glycan structures including *N-* and *O-*linked glycans and chitin, a polymer of GlcNAcβ1,4GlcNAc. Several lectins predominantly recognize GlcNAc termini, although the context of recognition is lectin specific. These include *Griffonia simplicifolia-II* (GS-II), *Phytolacca Americana* (PWA), *Ulex Europaeus-II* (UEA-II) and Wheat Germ Agglutinin (WGA) (**Figure 8**). A detailed analysis of these lectins is given below.

### Griffonia simplicifolia-II (GS-II)

At least four distinct lectins, GS-I to GS-IV, have been isolated from the seeds and leaves of *Griffonia simplicifolia*, also known as *Bandeiraea simplicifolia,* a shrub native to West Africa [93]. These include *Griffonia simplicifolia-I* (GS-I), discussed in the galactose section below, and *Griffonia simplicifolia-II* (GS-II), identified as a terminal GlcNAc binder [93, 94]. Machine learning confirmed terminal GlcNAcβ as the principal binding determinant of GS-II (**Figure 8**). The lectin prefers GlcNAc capped LacNAc in multiantennary *N-*glycans, or polyLacNAc with longer polymers. Chitin oligomers [(GlcNAcβ1,4)n, n≥2] are also bound. Terminal GlcNAcα is also recognized with multiple linkages tolerated (α1,2-,1,3-,1,4-,1,6-). Two different preparations of GS-II (EY, Vector) have almost identical binding patterns.

### Phytolacca Americana (PWA)

The pokeweed plant (*Phytolacca americana)* has at least six lectins, all of which are annotated as chitin binders [3]. Commercially available preparations of this lectin are a mixture of at least five of these proteins (E.Y. Laboratories). Machine learning analysis identifies chitin oligomers (GlcNAcβ1,4)n (n*≥*4), as the predominant binding motif for these lectins (**Figure 8**).

### Ulex Europaeus Agglutinin - II (UEA-II)

As previously discussed, the two main lectins identified from the seeds of the Furze gorse plants (*Ulex europaeus*) have distinct specificities. Unlike the fucose binding UEA-I, UEA-II is thought to bind β1,4-linked GlcNAc (i.e., chitin and related structures) [95]. Machine learning identified terminal GlcNAcβ1,3Galβ as the predominant binding motif (**Figure 8**). All glycans containing this motif were bound, however no terminal chitobiose motifs were recognized. Binding to oligosaccharides containing GlcNAcβ1,3GalNAc was also observed. In addition, minor binding to H-antigen motifs, also recognized by UEA-I, was also observed.

### Wheat Germ Agglutinin (WGA)

WGA, a lectin derived from wheat germ (*Triticum aestivum* or *Triticum vulgare*), is one of the most widely studied and commonly used lectins [96]. Although WGA is often annotated as a GlcNAc binding lectin, it is thought to have a “broad” specificity, binding sialic acids and a mixture of GlcNAc containing glycans [97]. We tested preparations from four vendors (EY, Vector, Seikagaku, Sigma) on CFGv5. Although preparation-specific differences were seen, machine learning indicates that the principal recognition motif for WGA is terminal GlcNAcβ (**Figure 8**). Presentation of this epitope on long chain polyLacNAc, multiantennary N-glycans or longer chitin oligomers enhanced binding. In keeping with literature, a wide variety of other terminal *N*-acetyl-containing glycans were recognized, including terminal GlcNAcα-, GalNAcα-, GalNAcβ-, MurNAcβ- and, in some preparations, Neu5Ac.

### Terminal Gal and LacNAc Binding Lectins

Terminal galactose is observed in a wide variety of contexts, from the immunogenic *α*-Gal epitope to the ubiquitous Galβ structures of Type I and Type II LacNAc commonly observed on *N-* and *O-*glycans and glycolipids [20]. The following lectins predominantly bind either terminal galactose or LacNAc epitopes: *Bauhinia purpurea* lectin (BPA, BPL)*, Erythrina Cristagalli* agglutinin (ECA, ECL), *Griffonia simplicifolia-I* (GS-I), *Lycopersicon Esculentum* agglutinin (LEA), *Marasmius oreades* agglutinin (MOA), *Pseudomonas aeruginosa-IL* (PA-IL), *Ricinus Communis* agglutinin (*RCA-I, RCA120*), *Sophora Japonica* agglutinin (SJA) and *Solanum Tuberosum* lectin (STA, STL) (**Figure 9**). The lectin from *Datura stramonium* (DSA) is also a LacNAc binder and is discussed earlier in this work as it is *N*-glycan specific (**Figure 4**). A detailed description of the binding specificities of these lectins follows.

### Bauhinia purpurea agglutinin (BPA, BPL)

Isolated from camels foot tree, *Bauhinia purpurea* agglutinin or lectin (BPA, BPL) is thought a GalNAc specific lectin [98]. Machine learning analysis identified the principal binding determinant as terminal β-Gal, with a preference for β1,3- over β1,4-linkages (**Figure 9**). The underlying residue can be GlcNAc (as in LacNAc) or GalNAc. This is one of the few lectins that shows a preference in binding for Type I LacNAc over Type II. BPL tolerates substitution on the internal GlcNAc of LacNAc, including fucose. In addition, BPL recognizes terminal β-GalNAc attached to either Gal or in Type I/II LacdiNAc motifs, tolerating fucosylation on the internal GlcNAc. However, binding is inhibited by α2,3- sialylation on proximal Gal residues resulting in no recognition of GalNAcβ-terminated glycosphingolipids, including GM2 (GalNAcβ1,4(Neu5Ac *α*2,3)Gal*β*1,4Glc), GD2, GT2 and related structures.

### Erythrina Cristagalli agglutinin (ECA, ECL)

Isolated from the seeds of the cockscomb plant, *Erythrina Cristagalli* agglutinin (ECA) was first identified as a Gal/GalNAc binding lectin [99]. It is reported that ECA bound exclusively to various terminal LacNAc structures, polyLacNAc, and branched *O*-glycans [100]. In line with this, machine learning identifies terminal Type 2 LacNAc as the predominant binding epitope (**Figure 9**). Binding is enhanced by the presentation of LacNAc in multiantennary contexts (*N-* or *O-*) or terminating polymers of Type 2 LacNAc (n≥2 repeats, i.e., a minimum of tetrasaccharide). This lectin also recognizes unsubstituted terminal Type 2 LacdiNAc (GalNAcβ1,4GlcNAc) containing glycans. Binding of Type 1 LacNAc or Type 1 LacdiNAc are not observed on CFGv5. In agreement with the previous literature [100], we also observe some weak binding to epitopes containing terminal Fucα1-2Galβ1,4GlcNAc.

### Griffonia simplicifolia-I (GS-I)

GS-I, from *Griffonia simplicifolia,* is a mixture of five isolectins, and is specific for α- galactosyl groups (Gal and GalNAc) [101, 102]. In concordance with the known specificity, machine learning identifies the principal binding motif for GS-I as terminal α-Gal (**Figure 9**). Machine learning also reveals inhibition of binding by α1,2-fucosylation on proximal residues, such as seen in blood group B. Glycans containing terminal GalNAcα, including the Tn antigen, are also bound.

### Lycopersicon Esculentum (LEA, LEL)

Isolated in 1980, the *Lycopersicon Esculentum* Agglutinin (LEA, LEL) from tomato plants binds GlcNAc oligomers, polyLacNAc and/or chitin [3, 103]. Analysis of LEA from two different commercial sources (EY, Vector) showed overlapping binding patterns. Machine learning identified chitin oligomers as the predominant binding motif for the EY preparation (**Figure 9**). In contrast, Type II polyLacNAc was identified as the predominant binding motif for the Vector preparation. Each preparation (EY, Vector) was found to also bind the predominant motif identified for the other preparation. LEA was permissive for various substitutions on LacNAc, including sialic acids (Neu5Ac or Neu5Gc, 5,9- diacetylated sialic and KDNa2), 3’- or 6’-sulfation, α1,2-fucosylation and terminal GlcNAc. Type II LacdiNAc was also bound. In contrast, Type I LacNAc was only recognized if the epitope was terminal on a Type II LacNAc core, arguing that this motif is not recognized by LEA.

### Marasmius oreades agglutinin (MOA)

Purified and cloned in 2002, the agglutinin from the mushroom *Marasmius oreades* (MOA) binds the xenotransplantation antigen Galα1,3Gal and blood group B [Galα1- 3(Fucα1-2)Gal] [104]. In agreement with the literature, machine learning analysis identifies Galα1,3Gal as the predominant binding motif (**Figure 9**). This motif lies within the B trisaccharide, which is preferentially bound by this lectin. MOA is sensitive to internal structures, as fucosylation of GlcNAc residues (as in Galα1-3Galb1-4(Fucα1-3)GlcNAc) inhibits binding. The closely related structure Galα1,3GalNAc is also recognized by MOA, in line with previous work [105].

### Pseudomonas aeruginosa-IL (PA-IL, LecA)

The bacterial lectin PA-IL, also known as LecA, from *Pseudomonas aeruginosa* exhibits affinity for α-galactosylated glycans [106, 107]. In keeping with this, machine learning analysis identified the predominant binding motif as terminal α-Gal (**Figure 9**). However, binding is inhibited by the presence of α1,2-fucosylation on the proximal residue, resulting in greatly diminished binding to blood group B antigens.

### Ricinus Communis agglutinin (RCA-I, RCA120)

Castor beans (*Ricinus Communis*) contain two similar, but distinct, lectins, the potent cytotoxin ricin and the substantially less toxic RCA-I (RCA120). Both lectins bind galactose and each is encoded by a unique gene [108, 109]. RCA-I from two commercial sources (EY, Vector) showed similar binding patterns. Machine learning identified terminal Type 2 LacNAc as the main binding determinant (**Figure 9**). RCA-I tolerates substitution at the 6-position of the terminal galactose, including sulfation and α2,6-sialylation, but not at the 3-position.

### Sophora Japonica agglutinin (SJA)

Isolated from the seeds of the Japanese pagoda tree, the *Sophora japonica* agglutinin (SJA) is known to bind both GalNAc and Gal (GalNAc > Gal), with an affinity for blood group B antigen [110, 111]. Machine learning identified terminal Type 2 LacNAc on multiantennary branches or polyLacNAc chains containing > 5 Gal residues as the predominant binding motif (**Figure 9**). This motif, however, only covered ∼1/3 of binders. Glycans terminated with either blood group B or type 2 LacdiNAc were also strongly bound, indicating that all three motifs are recognized by this lectin.

### Solanum Tuberosum (STA, STL)

*Solanum Tuberosum* Agglutinin (STA, STL), isolated from potato, is reported to be a polyLacNAc and chitin binder [3, 112]. Machine learning analysis identified internal linear Type 2 LacNAc as the major binding motif (**Figure 9**). Similar to LEA, STA tolerates a wide variety of termini, including sialic acid substituents. However, STA prefers linear glycans and binding is diminished by branching (bi-, tri- or tetraantennary), indicating that binding requires a free access to the linear chains. STA also binds chitin oligomers (n=2-5) and Type 2 LacdiNAc. Analysis of a second preparation of STA (EY) at a single concentration showed similar binding specificities.

### Terminal GalNAc binding lectins

Exposed GalNAc residues are carried by a wide variety of oligosaccharides, including blood group A [GalNAcα1-3(Fucα1-2)Gal], LacdiNAc, the Forssman antigen (GalNAcα1-3GalNAc) and the Tn antigen (GalNAcαSer/Thr). The following lectins predominantly recognize GalNAc termini and are discussed below (**Figure 10**): *Cytisus scoparius Lectin* (CSA), *Dolichos biflorus* (DBA), *Soybean agglutinin* (SBA)*, Vicia villosa lectin* (VVL, VVA), *and Wisteria floribunda* agglutinin. Tn antigen binding lectins from *Codium fragile* (CF)*, Helix aspersa* (HAA) and *Helix pomatia* (HPA) are discussed in **Figure 5**.

### Cytisus scoparius lectin (CSA)

Isolated from the Scotch broom plant *Cytisus scoparius,* the agglutinin CSA has been identified as a GalNAc specific lectin [113]. In keeping with this, machine learning analysis identified the predominant binder as terminal β-GalNAc (**Figure 10**). CSA shows a preference for LacdiNAc epitopes. In general, binding to terminal β-GalNAc containing glycans is inhibited by the presence of sialylation or fucosylation at the 3-position of proximal residues. For example, glycosphingolipids such as GM2 are not recognized. A limited subset of simple mono and disaccharide terminal α-GalNAc epitopes are also bound by this lectin.

### Dolichos biflorus agglutinin (DBA)

Isolated from the South Asian horse gram plant, *Dolichos biflorus* agglutinin (DBA) is used as a probe for terminal α-GalNAc residues and is used to bind blood group A [114]. Machine learning on the binding patterns of DBA from two sources (EY and Vector) unequivocally identified the Forssman antigen as the best binding motif (**Figure 10**). This is in line with previous reports that identified this antigen as a far stronger binding epitope than blood group A [115]. DBA also weakly bound to GM2 and related structures. Despite literature, no significant binding to blood group A was observed except at the highest concentration of lectin tested, where weak binding could be seen.

### Soybean (Glycine max) agglutinin (SBA)

The lectin from *Glycine max* seeds, otherwise known as soybeans, is known as a GalNAc binder [116, 117]. SBA from two different sources (Vector and EY) have consistent binding profiles. Machine learning identified both terminal β-GalNAc and the α-GalNAc terminated Forssman antigen as predominant motifs (**Figure 10**). Similar to CSA, the lectin prefers β-GalNAc in terminal LacdiNAc glycans and is sensitive to glycosylation at the 3-position of the proximal residue. Weaker binding to α-GalNAc- epitopes beyond the Forssman antigen is also observed, although fucosylation of the proximal residue, (e.g., as in blood group A) diminishes binding.

### Vicia villosa lectin (VVL, VVA)

The seeds of the hairy vetch plant, *Vicia villosa*, contain several lectins with distinct glycan binding specificities [3]. The lectin commonly annotated as VVL (or VVA) is a GalNAc binder [118, 119]. The preferred binding motifs of this lectin are almost identical to those of CSA. Machine learning on binding data of VVL from two sources (EY, Vector) identified both terminal β-GalNAc and LacdiNAc as predominant epitopes (**Figure 10**). Similar to CSA and SBA, binding is inhibited by fucosylation or sialylation of the proximal 3-position of β-GalNAc terminated glycans. Binding to a subset of simple mono and disaccharide terminal α-GalNAc epitopes (e.g., Tn), but not the more complex blood group A, is also observed, concordant with literature reports [119].

### Wisteria floribunda agglutinin (WFA, WFL)

The lectin from *Wisteria floribunda* (WFA, WFL) has been reported to recognize terminal GalNAc structures with high affinity, particularly those bearing LacdiNAc [120, 121]. In line with this, machine learning analysis identified its principal binding motif as terminal β-GalNAc (**Figure 10**). Unlike other lectins in this group (CSA, SBA, VVL), WFA is tolerant of substitution on the proximal residue and can bind β-GalNAc terminated glycosphingolipid structures such as GM2. This lectin also recognizes simple terminal α- GalNAc epitopes. WFA shows significant binding to multiantennary glycans bearing terminal LacNAc epitopes (both Type I and II), indicating that although terminal GalNAc is preferred, it is not absolutely required for binding.

## CONCLUSIONS

Lectins are a major tool for glycan analysis, finding use in flow cytometry, ELLA assays, lectin blots, histology and lectin microarrays. Despite their ubiquitous presence in glycosylation research, proper annotation of their specificities is still limited. Herein, we have used a mixed machine learning and manual annotation approach to analyze 116 lectin preparations, providing annotation for 57 unique lectins. These lectins cover the majority of those used in the literature. In general, our analysis found good concordance between preparations of the same lectin, regardless of source, with some lot to lot variation, which are most likely due to differences in the natural products. This points to the importance of cross-validating lectin results from naturally isolated lectins, for example through the use of an array with multiple probes. It also showcases the need for more recombinant lectins and antibodies [122–124].

Our analysis brings new insights into lectin specificities, finding both known binding motifs and previously unknown requirements for lectin binding. For example, MAA-I, commonly used to detect *α*2,3-sialic acid, was confirmed to preferentially bind 3’-O sulfation as previously reported. Machine learning identified that this lectin is inhibited by fucosylation of the proximal GlcNAc, as in sialyl or sulfo-Lewis x, a new finding. The more detailed annotation provided by this work sets the stage to garner additional insights from lectin binding. This includes more advanced annotation of motifs from lectin microarrays and other multi-lectin studies.

## MATERIALS AND METHODS

### Lectins and Antibodies

Biotinylated lectins were purchased from the following companies: E.Y. Laboratories (San Mateo, CA), Vector Laboratories (Burlingame, CA), Seikagaku Corporation (Tokyo, Japan), and Sigma (St. Louis, MO). Lectins that were not available in biotinylated form were biotinylated using the Pierce Biotinylation kit (Pierce, Rockford, IL) and standard methods. For a complete list of lot numbers and vendors for lectins please see **Supplemental Table S2**.

### Glycan Microarray Analysis

All GBPs were analyzed on the Consortium for Functional Glycomics Glycan microarray version 5.0 as detailed in [11]. The raw data is available at http://www.functionalglycomics.org/static/index.shtml.

### Data Analysis

We generated Z-score data for each glycan microarray, excluding those whose highest signal was <1000 RFU. For each GBP, we then generated a master Z-score (Zs) using Z-score data from multiple concentrations via Stouffer’s method [18]. In brief, the Z-scores for a glycan across concentrations were summed and divided by the square root of the number of concentrations. Calculations were done using Microsoft Excel 2011. Only GBPs for which multiple concentrations were tested and had more than a single array with signals > 1000 RFU were considered in our analysis. We used a threshold of Zs=1.645 (p≤0.05) to establish significance. Glycans that failed to meet this threshold for all GBPs tested were removed from our analysis (see **Table S1**). For each lectin, glycans that met our Zs threshold but did not vary across concentrations (σ^2^<10% of maximum variance across arrays) were set to average prior to generation of the hierarchical cluster and were flagged for careful consideration in our annotation. If these glycans gave signals of <10% of the maximum signal at the highest lectin concentration, they were considered non-specific binders in our motif analysis. Data was then annotated by hand to identify binding motifs. Heatmap was generated using Cluster 3.0 using the Pearson correlation coefficient as the distance metric and average linkage analysis and visualized with Java Treeview.

### Data Processing

For each glycan, we subtracted the average binding Z-score across all experiments to correct for unspecific background binding. We further removed several glycans from downstream analysis as their binding pattern indicated unspecific binding (**Table S1**).

### Hierarchical clustering to determine lectin binding specificity

Using the processed Zs data, we generated hierarchically-clustered heatmaps for all remaining glycans. For this, we removed several lectins that did not show a well-defined binding pattern (LPA_EY, NPA_EY, VRA_EY, BDA_EY, SJA_EY, GS-I_EY, HMA_EY, PWA_EY, TKA_EY, SVAM_Sigma, ACL_EY, LTL_EY). In addition, LcH_EY was clearly mislabeled as it was almost identical to AIA. Therefore, these lectins were removed from subsequent analysis. The heatmap was then generated using the Multiexperiment Viewer software (MeV_4_8, v.10.2). Samples were clustered heiarchically using the Pearson correlation as the distance metric and average linkage analysis.

### Extracting glycan-binding rules for lectins using machine learning

We used the Python package <u>SkopeRules</u> to identify logical, interpretable rules for lectin binding specificity. This procedure extracts rules from ensembles of tree-based machine learning models [125] that explain the glycan binding behavior of a given lectin using glycan features as input variables. The underlying machine learning models were trained on 80% of the processed data to predict whether a given glycan was bound (Z- score >= 1.645) in the remaining 20% of the data. Our ensembles consisted of 50 estimator models with a depth of one. As input features for these models, we used both 68 hand-annotated glycan features (refer to supplementary table) as well as the count of all observed mono- and disaccharide motifs and linkages. For each rule, we used the default thresholds recommended by SkopeRules of 0.3 for precision and 0.1 for recall to determine if a rule is valid. In this context, precision refers to the fraction of glycans fulfilling a given rule that bind to the lectin of interest while recall indicates how many of all bound glycans for that lectin fulfill the rule. Additionally, we only considered rules that had been chosen by the algorithm in at least 15% of the estimators to increase robustness. If no rule that fulfilled these requirements could be identified at a depth of one, we searched for rules among estimators with a maximum depth of two. If two rules were identified that both satisfied our quality criteria, we indicated them as alternative rules on the same level (e.g., rule 1a and 1b). After identifying the first rule, we selected all glycans fulfilling that rule and searched for another rule that improved the prediction of the glycan binding behavior. We repeated this procedure until we either could not identify another rule with our minimal requirements, reached a precision of 1.0, or reached five levels of rules.

## ACKNOWLEDGEMENTS

This work would not be possible without the work of the pioneers who came before including Els Van Damme and the late Profs. Irwin Goldstein and Hans-Joachim Gabius. This work was funded by the National Institutes of Health (Bridging Grant), the Canada Excellence Research Chairs Program (L.K.M.) and by the National Institute of Allergy and Infectious Diseases, a component of the NIH, Department of Health and Human Services, under contract 75N93019C00052.

