## Supplemental Figures for "A Useful Guide to Lectin Binding: Machine-Learning Directed Annotation of 57 Unique Lectin Specificities"

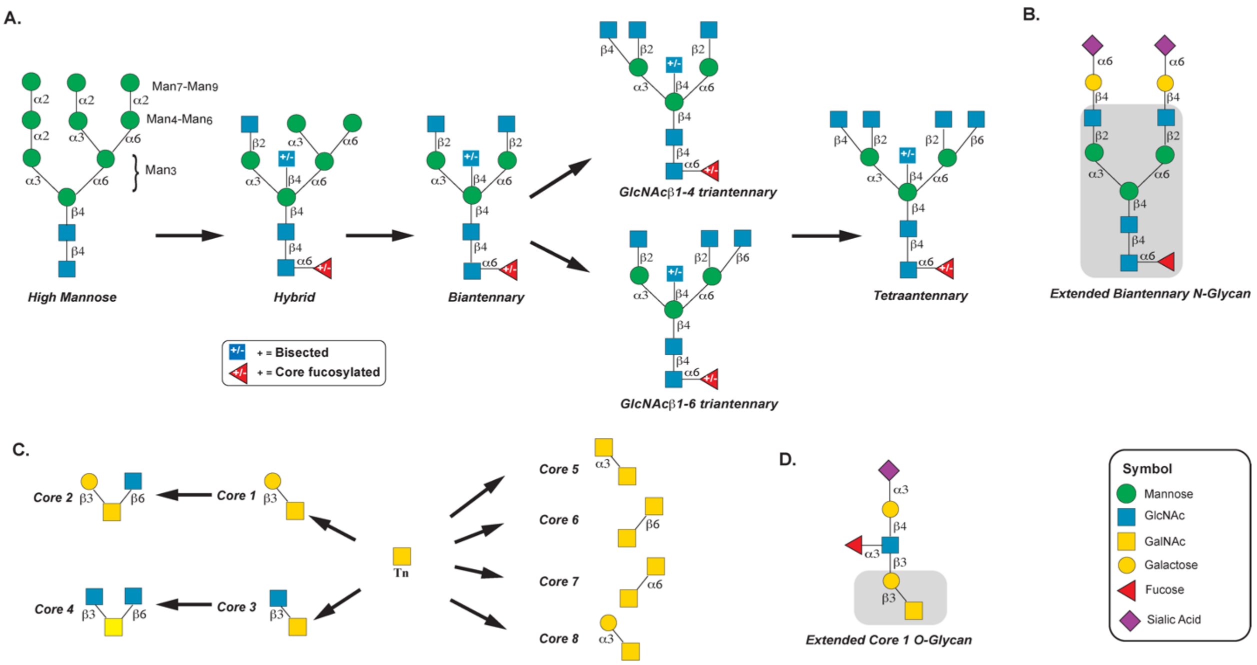


**Figure S1.** Overview of *N*- and *O*- glycans. A. *N*-glycosylation pathway. High mannose is trimmed $Man_5$ and then elaborated to hybrid, biantennary, triantennary and tetraantennary *N*-glycans. B. An example of a more complex biantennary *N*-glycan. The biantennary core is highlighted in grey. C. *O*-glycan cores. D. An example of a more complex core 1 O-glycan. The core is highlighted in grey.

**Tables S1-5:** Excel sheets.
